# The cerebrospinal fluid virome in people with HIV: links to neuroinflammation and cognition

**DOI:** 10.1101/2025.02.28.640732

**Authors:** Mattia Trunfio, Rossana Scutari, Valeria Fox, Elisa Vuaran, Raha Maryam Dastgheyb, Vanessa Fini, Annarita Granaglia, Francesca Balbo, Dora Tortarolo, Stefano Bonora, Carlo Federico Perno, Giovanni Di Perri, Claudia Alteri, Andrea Calcagno

## Abstract

Despite effective HIV suppression, neuroinflammation and neurocognitive issues are prevalent in people with HIV (PWH) yet poorly understood. HIV infection alters the human virome, and virome perturbations have been linked to neurocognitive issues in people without HIV. Once thought to be sterile, the cerebrospinal fluid (CSF) hosts a recently discovered virome, presenting an unexplored avenue for understanding brain and mental health in PWH. This cross-sectional study analyzed 85 CSF samples (74 from PWH on suppressive antiretroviral therapy, and 11 from controls without HIV, CWH) through shotgun metagenomics for DNA/RNA viruses. Taxonomic composition (reads and contigs), α and β diversity, and relative abundance (RA) of prokaryotic (PV), human eukaryotic (hEV), and non-human eukaryotic viruses (nhEV) were evaluated in relation to HIV infection, markers of neuroinflammation and neurodegeneration, cognitive functions, and depressive symptoms. Sensitivity analyses and post-hoc cluster analysis on the RA of viral groups and blood-brain barrier permeability were also performed.

Of 46 read-positive CSF samples, 93.5% contained PV sequences, 47.8% hEV, and 45.6% nhEV. Alpha diversity was lower in PWH versus CWH, although p>0.05. At β diversity analysis, HIV status explained 3.3% of the variation in viral composition (p=0.016). Contigs retained 13 samples positive for 8 hEV, 2 nhEV, and 6 PV. Higher RA of PV was correlated with higher CSF S100β (p=0.002) and β-Amyloid 1-42 fragment (βA-42, p=0.026), while higher RA of nhEV with poorer cognitive performance (p=0.022). Conversely, higher RA of hEV correlated with better cognition (p=0.003) and lower βA-42 (p=0.012). Sensitivity analyses in virome-positive samples only confirmed these findings. Three CSF clusters were identified and showed differences in astrocytosis, βA-42, tau protein, and cognitive functions. Participants with hEV-enriched CSF showed better cognitive performance compared to those with virus-devoid and nhEV-enriched CSF (models’p<0.05).

This study provides the first comprehensive description of the CSF virome in PWH, revealing associations with neuroinflammation and cognition. These findings highlight the potential involvement of the CSF virome in brain health and inform about its composition, origin, and potential clinical implications in people with and without HIV.

**Author Summary:** HIV can affect brain health and mental well-being, even in people on successful antiretroviral therapy. The reasons behind this are still unclear. HIV also influences the communities of microbes and viruses living in the human body, and recent research suggests that the human virome, the collection of all viruses within the body, may play a role in cognitive functions, mood, and brain health. For a long time, scientists believed that the cerebrospinal fluid (CSF), which surrounds the brain, was sterile, while robust evidence has shown that the CSF hosts its own unique virome. Via advanced genetic sequencing (shotgun metagenomics), we analyzed the CSF virome in people with and without HIV looking for possible links to neuroinflammation, cognitive performance, and depression. We found that while HIV infection does affect the composition of CSF viral communities, there were no remarkable differences in the CSF virome of individuals with and without HIV. Most viral sequences appeared to come outside the brain. A higher abundance of non-human viral sequences, such as viruses of bacteria, plants, fungi, and animals, was associated with neuroinflammation and poorer cognitive performance. On the other hand, a higher abundance of human viruses correlated with better cognitive function and healthier signature of neuromarkers. These findings provide new insights into the presence and characteristics of the human CSF virome and how it might influence brain health. They also suggest new potential mechanisms of HIV-associated neuropathology.

## Introduction

The recognition of the human virome, is not recent, even for the “sterile” central nervous system (CNS). Viruses that establish latent chronic infections have long been recognized as permanent residents of the human body, and neurotropic viruses (e.g., *Herpesviridae*, *Retroviridae*) have been represented the CNS virome [1]. Thanks to next-generation sequencing, the human virome has expanded to include a broader range of non-pathogenic viruses persistently or transiently colonizing the body [2]. It is now recognized that the largest part of the human virome consists of commensal viruses, primarily bacteriophages (phages), that infect non-human cells [2–5].

The human virome is not a passive passenger: viral communities across the body exert local and systemic effects by which they can also influence distant organs such as the CNS [2,6–9]. Gut phages influence the host’s microbiome, metabolism [10], immune system [7,11], and disease risks [12,13]. The gut phages *Siphoviridae* and *Microviridae* affect cognition and behavior in humans, mice, and *Drosophila*, with their fecal transplant and oral supplementation promoting memory-related gene expression [9]. In mouse models of chronic stress, a *Microviridae*-enriched virome alleviates anxiety and restores stress-induced changes in circulating immune cells, cytokines, and gene expression in the amygdala [14]. *Chlorovirus* ATCV-1 in the human oropharynx is linked to lower cognitive performance, and can alter synaptic plasticity and learning/memory-related genes in the hippocampus [15].

While research is uncovering the effects of the gut microbiome on human brain and mental health, our understanding of the role of the CNS virome in maintaining or disrupting its host organ’s health is limited. For long, few bacterial, parasitic, and viral infections have been the target of investigations of a microbial etiology of neuroinflammation and CNS disorders [1,16–19], and a comprehensive study of the human CNS virome has just begun. As for other body sites, a rich variety of prokaryotic, non-human, and human eukaryotic viruses with and without pathogenic potential or tropism to the CNS have been detected in human brain and cerebrospinal fluid (CSF) in the last decade [20–25]. Yet, only one study assessed the role of these CNS viral communities in mental health, reporting different richness and composition of the virome in postmortem Brodmann Area 46 tissues between people suffering from schizophrenia and healthy controls [23].

HIV impacts the human microbiome throughout all stages of infection [26,27]. HIV-induced changes in immune cell activation, senescence, and exhaustion lead to shifts in the microbial communities, including viruses, across the human body [26,28–31]. While the dynamics and causality in the relationships between HIV, microbiomes, and host warrant further understanding [32], current evidence suggests a vicious cycle where microbiome changes following HIV-associated inflammation and organ injury can contribute to further systemic inflammation and increased risk of comorbidities [26,28,33,34].

HIV is also a neurotropic virus that impacts brain and mental health. Aging PWH face up to 58% greater risk of all-type dementia compared to age-matched people without HIV [35,36]. The global prevalence of HIV-associated cognitive disorders is estimated around 40% [37,38], and depressive disorders are two to three times more common in PWH than in the general population [39]. Neuroinflammation plays a pivotal role in both cognitive and mood issues in PWH [40,41], but the mechanisms and causes remain poorly understood, especially during suppressive ART. In people without HIV, the gut-brain microbiome axis is an emerging player in neuropsychiatric comorbidities [42–45], alongside several viral co-infections [46–52]. Similarly, immune responses against CMV, EBV, and HSV-1/2 have been associated with neuroinflammation, neurodegeneration, and poorer cognitive performance in PWH [53–55]. Differences in the virome in postmortem prefrontal cortex tissue have been described between people with and without HIV [25]. However, the characteristics and role of the CNS/CSF viromes have yet to be described.

In this pilot study we investigate the presence and composition of the CSF virome in PWH. We hypothesized that the CSF virome of PWH differs in diversity and composition from that of people without HIV. We expected that its characteristics are associated with factors that are both site-specific (e.g., blood-brain barrier -BBB-permeability) and common to other viromes of the human body (e.g., CD4+ T cell count). We further hypothesized that these differences would be associated with distinct levels of neuroinflammation and thereby with different degrees of cognitive performance and depressive symptoms. To evaluate these hypotheses, we performed shotgun viral metagenomics in CSF samples from PWH and control participants without HIV infection (CWH), and we assessed the virome in relation to HIV status, neuroinflammation, neurodegeneration, BBB permeability, cognitive performance, and depressive symptoms. The CSF virome was described using both an assembly-free (reads) and an assembly-based taxonomic annotation (contigs) after two considerations: 1) short viral genetic fragments from degraded virions can trigger immune responses through DNA/RNA sensing and other mechanisms [56,57]; these short immunologically active sequences can be discarded by contig assembly, underestimating the quantification of biological players [58]; 2) The choice of using one method over the other should be based also on the characteristics of the target virome (e.g., expected richness, occurrence of rare species, average length and abundance of reads), and very limited data are available on the human CSF virome to tailor a pipeline on established references. Through this hypothesis-generating study we compared these two approaches to provide further information to future studies in the field.

Consistent with our main hypothesis, we found a mild but significant divergence in the composition of the CSF virome by HIV status, and that variations in the relative abundance of human and non-human viruses in the CSF of PWH are associated with distinct signatures of neurodegenerative biomarkers and cognitive performance.

## Results

### 1. Study population

Eighty-five CSF samples from 74 PWH and 11 CWH were analyzed. As detailed in S1 Table, PWH were predominantly white (93.2%), male (67.6%), of middle age, with a long history of HIV infection and AIDS (62.2%). The routes of HIV transmission were equally represented. HIV RNA was detectable in the blood and CSF of 17.6% (range: 23-125 cp/mL) and 27.0% PWH (range: 20-133 cp/mL), respectively. Median CD4+ T cell count and CD4/CD8 ratio were 447 cells/µL and 0.7. CWH were also predominantly white, males, and mildly older than PWH (p=0.074; S1 Table).

## 2. Taxonomic2 composition of the CSF virome and differences by HIV status

Our primary aim was to describe the presence, taxonomic composition, and characteristics of the virome in the CSF of PWH and to compare it to that of CWH. Sequencing returned a total of 1487.78 million reads of mean length of 149 nucleotides and a median of 14.08 (9.87-21.81) million reads per subject. After quality filtering and removal of human-host reads, a total of 125.50 million non-human reads for 1.15 (0.54-1.90) million per subject were retained. After removing non-viral and blanks’ reads (detailed in S2 Table), 16,039 viral reads for 44 (12-131) reads per subject were retained. Eventually, taxonomy assignment (at least two unique reads and a confidence score greater than 0.5) retained 14,046 total reads for 46 CSF samples (54.1%; 17 [6–88] reads per sample), while contigs assembly retained 3,530 total reads for 13 CSF samples (15.3%; 142 [59–354] reads per sample).

### 2.1 CSF Virome according to Reads

Of the 46 samples positive for viral DNA or RNA, 38 were from PWH (51.3%) and 8 from CWH (72.7%). The higher prevalence of positive samples in CWH vs PWH did not reach statistical significance (Fisher’s p=0.214) even after adjusting by age that differed between the groups (aOR=2.95 [0.68-12.76] for CWH vs PWH, p=0.147).

Prokaryotic viruses (all phages, PV) comprised 71.2% of all CSF viral reads and were detected in 93.5% of the positive samples (median 30 reads [5–86] per sample), as detailed in Table 1.

**Table 1.**
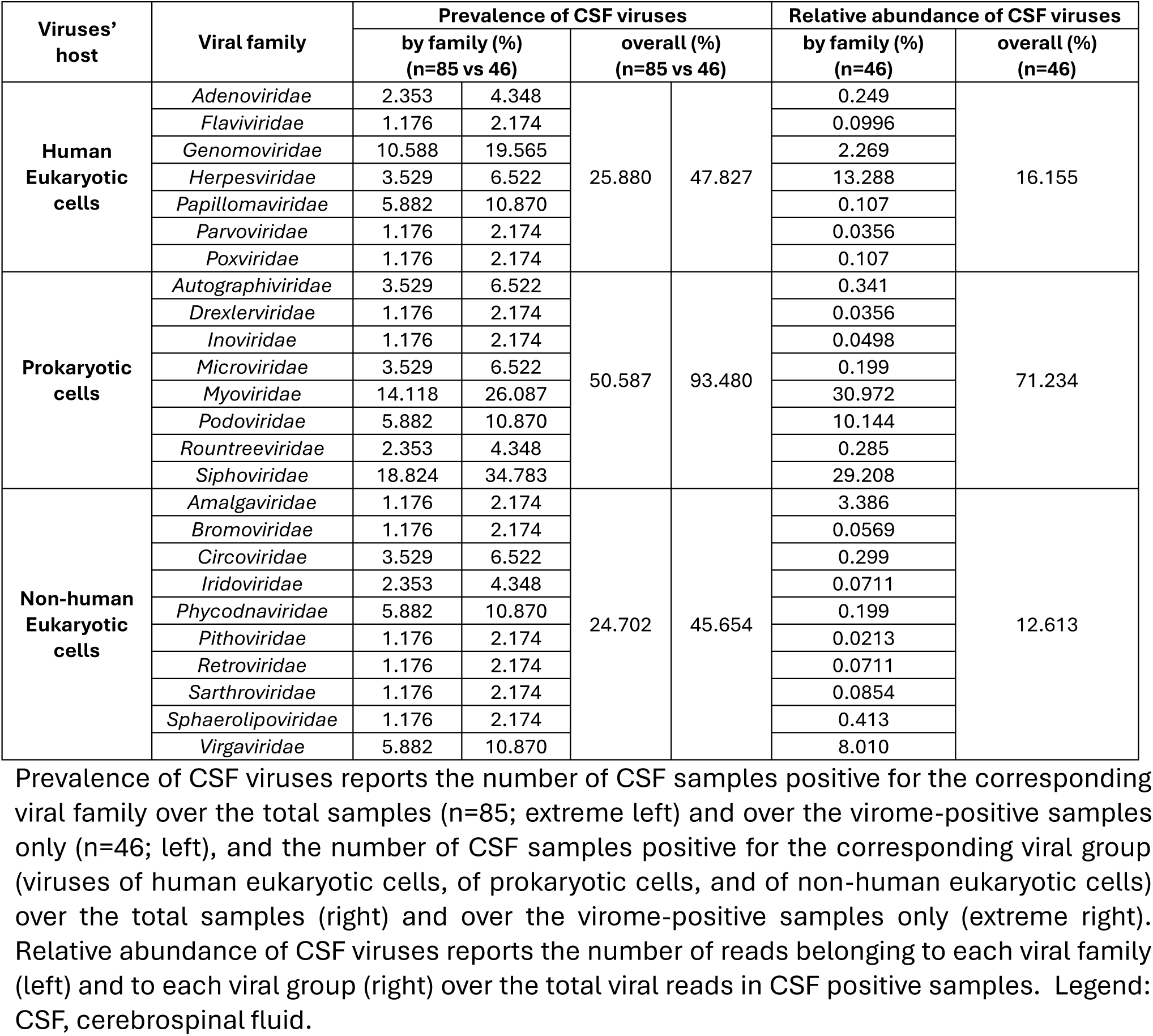
Taxonomic composition of the CSF virome according to reads.

Three families of the order *Caudovirales* (double-stranded DNA viruses), *Siphoviridae*, *Myoviridae,* and *Podoviridae* were the most abundant PV, being detected in 34.8%, 26.1%, and 10.9% of positive samples, and representing the 29.2%, 31.0%, and 10.1% of all the reads, respectively (Table 1).

Human eukaryotic viruses (hEV) made up 16.2% of the total reads and were detected in 47.8% of the positive samples (median 10 reads [4–33] per sample). Among these, *Genomoviridae* (single-stranded DNA) and *Papillomaviridae* (double-stranded DNA) were present in the 19.6% and 10.9% of virome-positive samples and accounted for the 2.3% and 0.11% of the total viral reads detected. *Herpesviridae* family (double-stranded DNA) followed by frequency (6.5%) but was the primary contributor to the CSF hEV reads (relative abundance, RA, 13.3%) (Table 1).

Non-human eukaryotic viruses (nhEV) constituted 12.6% of all CSF reads and were detected in 45.6% of the positive samples (median 4 reads [2–12] per sample). *Virgaviridae* (positive-strand RNA viruses of plants), *Phycodnaviridae* (double-stranded DNA viruses of algae)*, and Circoviridae* (single-stranded DNA viruses of animals) were the most frequent families. While *Virgaviridae* was also the main contributor to the total nhEV reads (8.0%), *Amalgaviridae* (double-stranded RNA viruses of plants and fungi) represented the second main contributor to nhEV reads (3.4%; Table 1).

The composition of CSF virome by viral family and their relative abundance for PWH and CWH is shown in Figure 1.

**Figure 1.**
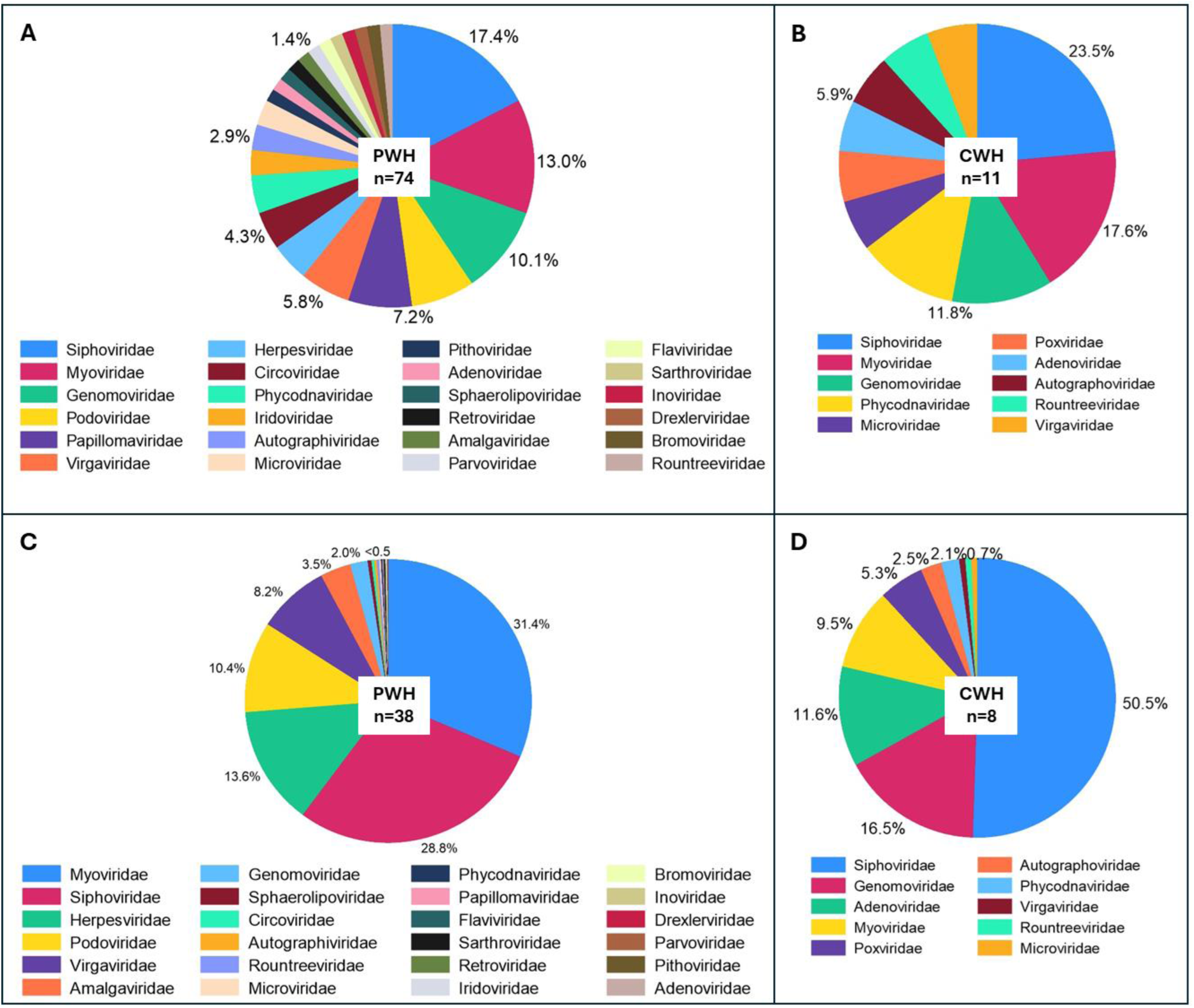
Prevalence and relative abundance of CSF viral families by HIV status Panels A and. **B** show the prevalence of viral families in the CSF (proportion of samples positive for each viral taxon among all the CSF samples) of PWH and CWH, respectively. Three families were found in about half of CSF samples from both PWH and CWH: *Siphoviridae* and *Myoviridae* (double-stranded DNA phages belonging to the order of *Caudovirales*), and *Genomoviridae* (single-stranded DNA viruses). **Panels C and D** show the relative abundance (number of reads of the specific taxon over the total number of reads per sample) of the viral families in virome-positive samples from PWH and CWH, respectively. *Myoviridae* and *Siphoviridae* represented the 60.2% of the total viral genetic material detected in the CSF of PWH, followed by *Herpesviridae* (13.6%, human double-stranded DNA viruses), *Podoviridae* (10.4%, double-stranded DNA phages belonging to the order of *Caudovirales*), *Virgaviridae* (single-stranded RNA viruses of plants), and 18 other viral families at much lower proportions. *Siphoviridae* contributed to the 50.5% of the total viral genetic material detected in the CSF of CWH, followed by *Genomoviridae* (16.5%), *Adenoviridae* (11.6%, human double-stranded DNA viruses), *Myoviridae* (9.5%), *Poxviridae* (5.3%, human double-stranded DNA viruses), and 5 other viral families at much lower proportions.

Alpha diversity was lower in samples from PWH compared to CWH: the number of observed taxa was 1.5 [0.0-2.0] vs 2.0 [1.0-2.0] and Shannon index was 0.00 [0-0.36] vs 0.32 [0-0.47], suggesting lower richness and less even distribution among taxa; Simpson index was 0.00 [0-0.14] vs 0.17 [0-0.30], confirming the dominance of fewer taxa in the CSF of PWH. Although all indexes were consistently lower in PWH, the differences did not reach statistical significance (p>0.05 for all, n=38 vs n=8; Figure 2.A). Instead, β diversity showed a significant divergence in the composition of CSF viral communities between PWH and CWH (p=0.016 for both the distance measures; Figure 2.B); however, HIV status explained only the 3.3% and 3.4% (R2 of Jaccard and Bray Curtis distance, respectively) of the variation in viral composition. When the RA of each viral family and viral metrics (RA of PV, hEV, nhEV) were compared by HIV status no difference was detected (Figure 2.C).

**Figure 2.**
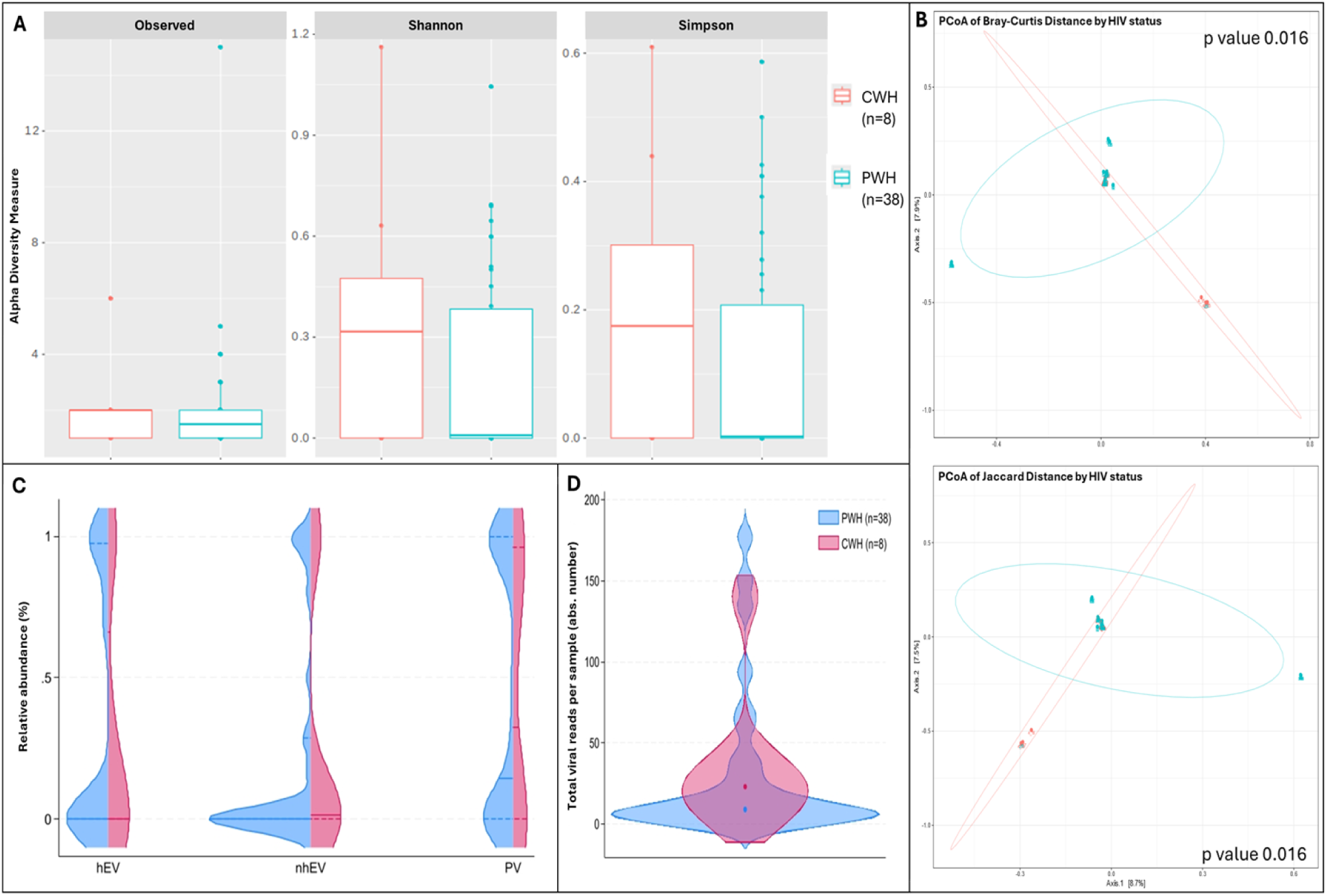
Alpha and Beta diversity and CSF viral metrics by HIV status Panel. **A** shows the number of observed viral taxa (left), Shannon index (middle), and Simpson index (right) of the CSF virome against HIV status (CWH in orange, PWH in blue). While CSF samples from PWH had lower median values for each of the three metrics, none reached statistical significance. **Panel B** shows the representation of β diversity based on Bray Curtis (upper) and Jaccard distances (lower; CWH in green, n=8; PWH in blue, n=38; each dot represents a sample). A significant divergence in the composition of CSF viral communities based on HIV status was observed; Permutation Based Analysis of Variance (PERMANOVA) with Adonis function. Violin plots in **Panel C** show the relative abundance of human eukaryotic viruses (hEV), non-human eukaryotic viruses (nhEV), and prokaryotic viruses (PV) reads in CSF samples from PWH (blue half; n=38) and CWH (red half; n=8); median and interquartile range are represented by continuous and dotted lines: hEV 0% (0-96.2) vs 0% (0-49.1), nhEV 0% (0-25.6) vs 1.4% (0-30.8), and PV 14.5% (0-100) vs 32.5% (0-94.2) in PWH vs CWH; when compared (Mann-Whitney U test), no significant difference was detected (p=0.782 for hEV, p=0.146 for nhEV, and p=0.641 for PV). **Panel D** shows the violin plots (overlap) of the total number of viral reads detected in the CSF of PWH (median 15 reads per sample [6–124]; n=38; blue violin) and of CWH (median 23 reads per sample [14–36]; n=8; red violin); the median is represented by the dot; when compared by Mann-Whitney U test, no significance was detected (p=0.247).

Sequencing did not identify HIV-1 (including the 20 CSF samples with PCR-positive HIV RNA), EBV (in all the 3 PCR-positive samples), and JCV (in both the PCR-positive samples). The detection limit of viruses in CSF through metagenomics, including HIV-1 RNA, has been previously described at ∼10^2^ copies/mL [59–61]; consistently, the viral load of these viruses in all the PCR-positive samples was <150 cp/mL (see S3 Table for details).

### 2.2 CSF Virome after Contigs reconstruction

When only viral reads assembled into contigs of at least 300 nucleotides were considered, 10,516 reads had insufficient length, coverage, and overlap. After filtering these out, we retained 13 CSF samples positive for virome: 10 from PWH (13.5%) and 3 from CWH (27.3%). Accordingly, alpha diversity was lower: in PWH, the number of observed taxa was 1 [1–1], and both Shannon and Simpson index 0.0 [0.0-0.0]; in CWH, the number of observed taxa was 1, 1, and 4, Shannon index was 0.0, 0.0, and 0.92, and Simpson index was 0.0, 0.0, and 0.48.

Contigs belonged to 8 unique hEV, 2 nhEV, and 6 unique PV (Figure 3). Among hEV, the dsDNA *Herpesviridae* family was represented by EBV and HHV-6 in one participant each, with 1,467 and 376 reads (genome coverage of 2.64% and 8.85%, respectively). HPV-12 and HPV-96 (in one participant each) were detected with 23 and 26 reads (genome coverage of 5.68% and 8.62%). The eukaryotic DNA viruses *Poxviridae Molluscum Contagiosum virus*, the *Human Mastadenovirus C* and the *Gemycircularvirus HV-GcV2* were also detected in one CWH each (27, 23 and 64 reads; genome coverage of 0.16%, 1.78%, and 33.7%). HCV was the only eukaryotic RNA virus identified and found in one sample from PWH with 98 reads and a genome coverage of 8.03%. The 2 nhEV were both viruses affecting tomato plants and other solanaceous crops, *Tomato Brown Rugose Fruit Virus* and *Southern Tomato Virus*, both found in one PWH, and presenting the highest genome coverage (98.3% and 94.6% respectively).

**Figure 3.**
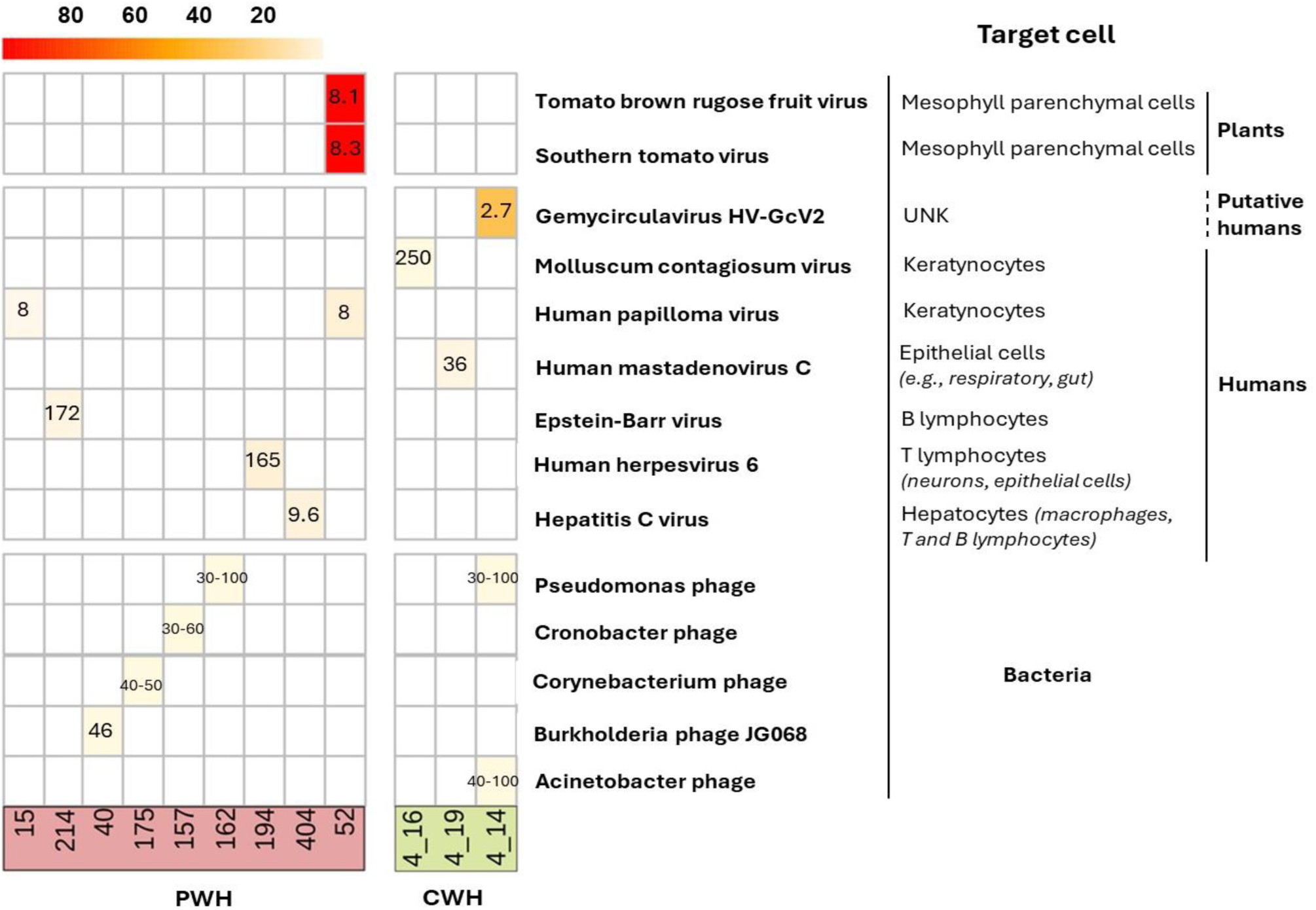
Heatmap of the genome coverage of CSF viruses detected after contig assembly. Each column corresponds to a CSF sample (PWH in red; CWH in green) and each row corresponds to the virus recovered. As shown in the heatmap, the genome coverage ranged from ∼5% for the bacteriophages to almost the 100% for two non-human eukaryotic viruses, *Tomato brown rugose fruit virus* and *Southern tomato virus*, both detected in the CSF of one participant with HIV infection (ID52). The length of the genome of each identified CSF virus is reported in the corresponding box (range min-max for taxa with variability among species and strains). As detailed on the right, thirteen out of sixteen (76.5%) contigs belonged to viruses that have no target cells residing in or transiting through the CNS; EBV, HHV-6, and HCV have no primary target cells within the CNS. The mean and median length of CSF viral genomes were 914 (±3859) and 2835 (750-2110) base pairs.

Of the 5 CSF samples positive for the 6 PV, all but one were from PWH (see S5 Table and S1 Material). None of these participants had a medical history indicating infections by the bacterial hosts of the identified phages (e.g., *Acinetobacter spp*, *Pseudomonas spp*). Of interest, the target hosts are all acknowledged commensal bacteria or opportunistic pathogens in the human gut, skin, and lung bacteriomes. The detailed characterization of these 12 participants is reported in S4 and S5 Table and S1 Material.

#### 3. Exploratory analyses of other factors associated with the CSF Virome

To assess whether other parameters were associated with the characteristics of the CSF virome, we performed exploratory analyses of the association between demographics and HIV-related parameters, α and β diversity, and viral metrics. Age, sex, demographics, and HIV-related parameters listed in S1 Table (e.g., CD4+ T cell count) were not associated with α diversity, viral metrics, nor with significant divergence of viral communities at β diversity analysis (data not shown). Due to the small number of positives, we did not perform these analyses after contig assembly.

### 3.1 CSF Virome and Neuroinflammation

As secondary aim, we assessed whether α diversity and viral metrics (RA of hEV, nhEV, PV) were associated with CSF biomarkers of neuroinflammation, neurodegeneration, and BBB permeability. We hypothesized that, regardless of the origin and size of the genetic material, its presence in the CSF can trigger immune activation and CNS injury. The CSF biomarkers are shown in S6 Table. The prevalence of BBB impairment was similar between PWH and CWH (21.6% and 18.2%), while intrathecal synthesis was detected only in PWH (27.0% vs 0%; S6 Table). CSF-to-serum albumin ratio (CSAR), intrathecal synthesis, and CSF biochemistry (glucose, leukocytes, and proteins) were not correlated with diversity and viral metrics (data not shown).

As for the biomarkers of neuroinflammation/neurodegeneration, the CSF levels of fragment 1-42 of β amyloid (βA-42) and S100β increased as the RA of PV increased (rho=0.366, p=0.002 and rho=0.267, p=0.026, respectively), while CSF βA-42 decreased as the RA of hEV increased (rho= -0.297, p=0.012). At sensitivity analyses performed only in virome-positive samples (n=38), the same correlations between CSF βA-42 and RA of PV and hEV were observed with larger effect size (rho=0.633, p<0.001 and rho= -0.435, p=0.008).

β diversity analysis did not show divergence in the composition of the CSF virome according to groups categorized by either the median value of the biomarkers in the study population or standard clinical cut-offs of the biomarkers (e.g., CSF proteins >45 mg/dL or CSF leukocytes >5 cells/mL; data not shown; the cut-offs of each biomarker are detailed in the Methods). No differences in α diversity, total number of viral reads, or viral metrics were also observed when participants, either all or only PWH, were compared by BBB impairment and presence of intrathecal synthesis (data not shown).

In summary, weak-to-moderate associations were observed between the RA of CSF phages and hEV and CSF biomarkers of amyloid metabolism and astrocytosis. Contrary to our expectation, impairment of the BBB and intrathecal synthesis were not associated with more viral genetic material in the CSF nor with distinct virome signatures.

### 3.2 CSF Virome, Cognitive performance, and Depressive symptoms

Then, we assessed whether CSF diversity and viral metrics were associated with cognitive performance and depressive symptoms. These analyses were performed in a subgroup of PWH with available data: 61 (82.4%) underwent neurocognitive evaluation, and 66 (89.2%) were evaluated for depressive mood. The prevalence of cognitive impairment was 34.4%, and depressive mood was also common (28.8%) but mostly mild (S6 Table).

Alpha diversity was not associated with global cognitive performance (Global Deficit Score, GDS), Domain Deficit Scores, nor depressive symptoms. Similarly, β diversity analysis showed no divergence in viral composition by depression (cut-off of BDI-II ≥14) and cognitive impairment (GDS ≥0.5), and the amount of CSF viral reads was also not associated with cognitive scores and depressive symptoms (data not shown).

On the contrary, viral metrics showed variable associations with cognitive performance. Specifically, higher RA of hEV was associated with better global cognitive performance (lower GDS; Figure 4), and with better scores at memory, attention/working memory, and executive functions (a trend for better verbal and motor functions was also observed; Figure 4). Higher RA of nhEV was associated with worse global performance and worse scores at executive functions (a trend for worse memory and motor functions was also observed; Figure 4). Higher RA of PV was borderline associated with worse global performance and executive functions. Depressive symptoms were not associated with viral metrics (Figure 4). At sensitivity analyses performed only in participants with virome-positive samples, the same correlations were observed (S7 Table).

**Figure 4.**
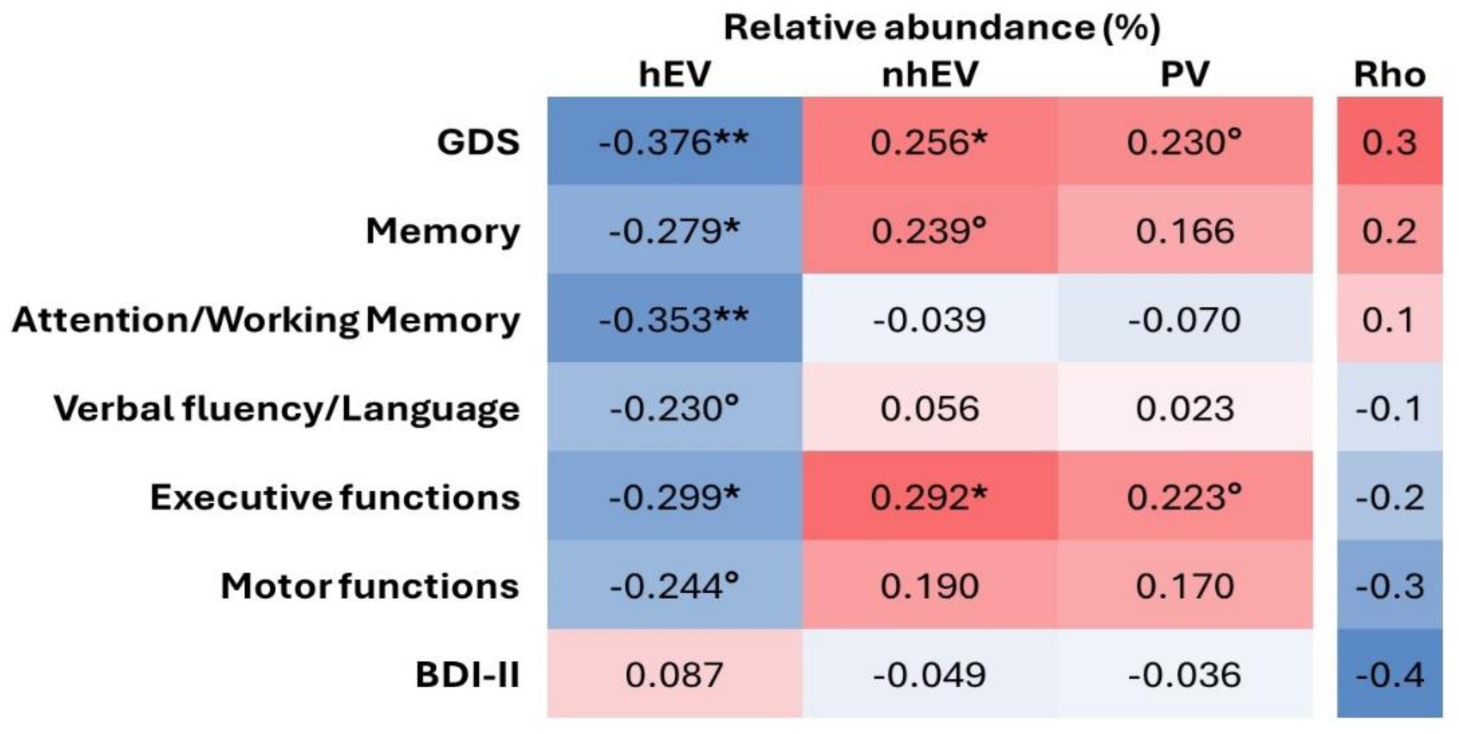
Heatmap of the correlations between relative abundance of CSF viral categories and neurocognitive measures. **p<0.005; *p<0.05; °p<0.1 in n=61 for cognitive metrics and n=66 for BDI-II. Legend: hEV, human eukaryotic viruses; nhEV, non-human eukaryotic viruses; PV, prokaryotic viruses; GDS, Global Deficit Score; BDI-II, Beck Depression Inventory II.

In summary, diversity and composition of the CSF virome were not associated with cognition nor depression, while the relative abundance of PV, hEV, and nhEV was differentially associated with global and domain-specific measures of cognitive performance.

#### 4. Post hoc analysis: CSF Virome Clusters

Having observed that a large proportion of the viral genetic material in the CSF belongs to viruses that do not infect cells residing in the CNS, we hypothesized that most of the genetic material detected in the CSF is constituted by genetic fragments coming from the periphery. Based on this hypothesis model, detailed in Figure 5A, we merged PV and nhEV into a single clustering variable (NHV) due to the similar nature (genetic fragments from peripheral sites) and activity (e.g., neuroinflammation through genetic sensing and activation of immune and scavenger cells [56,57]), and we clustered participants by the relative abundance of human and non-human CSF viruses (NHV) and BBB permeability. Our previous analyses explored associations assuming linear relationships and differences by demographics, cognition, and biomarkers, due to the lack of references to group participants according to the characteristics of the CSF virome. This post hoc analysis investigated the same relationships after clustering participants by CSF virome characteristics. As we explored associations between the CSF virome and mental health, we performed the clustering in PWH only. However, when CWH were included to assess the consistency of the clustering solutions, similar clustering results were observed (data not shown).

**Figure 5.**
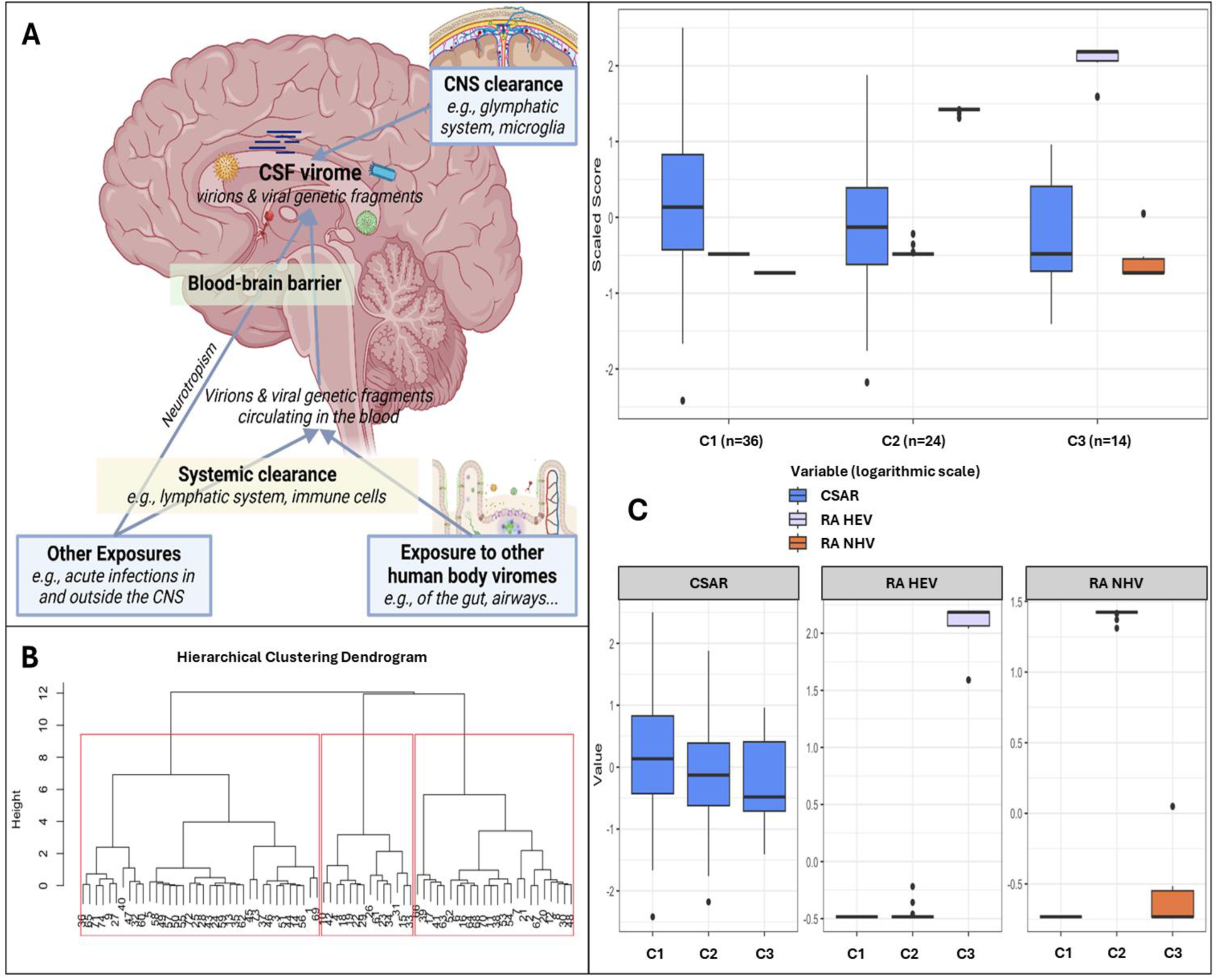
Clusters based on relative abundance of human and non-human CSF viruses and blood-brain barrier permeability Panel. **A** shows the hypothesis model: the human CSF virome is the collection of viral genetic material that can come from both live virions and fragments of genetic material from death viruses. Part of the virome is generated within the CNS from viruses infecting resident or transiting cells (e.g., neurotropic viruses, viruses transported into the CNS by cells through trojan horse mechanisms), while part is represented by viral fragments that escape immune clearance and that originate from viruses that infect cells in blood and in other peripheral sites (e.g., gut, airways). These fragments eventually can enter the CNS as either free fragments (e.g., at the choroid plexus, through transcytosis, pinocytosis, or at leaking points along impaired BBB) or through trojan horse mechanisms (e.g., cells that phagocyted the fragment in the bloodstream and then migrated into the CNS). Therefore, the CSF virome (amount and type of viral genetic material) will depend on a broad range of factors: e.g., spillover from the viromes of other body sites, ongoing viral infections, BBB permeability, peripheral immune clearance, and the clearance in the CNS operated by local cells and the glymphatic system. Within our study, the peripheral exposure, the systemic and the CNS clearance were unmeasured variables. **Panel B** shows the dendrogram for hierarchical clustering and the 3 clusters identified in red squares. **Panel C** shows the three clusters based on CSF-to-serum albumin ratio and relative abundance of human and non-human (prokaryotic and eukaryotic) viruses.

Three clusters were identified (Figure 5B-C): C1 (n=36) included participants with no viral reads in their CSF, despite the highest prevalence of BBB impairment (Figure 5C and S8 Table); compared to C1, C2 (n=24) had similar degree of BBB impairment, and the CSF was enriched in NHV and poor of hEV; C3 (n=14) had the lowest prevalence of BBB permeability, and the CSF was enriched in hEV and poor of NHV (Fig.5C). Figure 6 and S8 Table show the comparisons of demographics, HIV-related parameters, biomarkers, and neurocognitive metrics between clusters. Among the differences, C2 had the highest prevalence of female sex, and levels of CD4+ T cells and astrocytosis (S100β protein). C3 had the lowest levels of all CSF biomarkers, with total tau, Aβ42, and leukocytes reaching statistical significance. None of the members of C3 had cognitive impairment (versus 56% in C2, p=0.005, and 32% in C1, p=0.048), attaining the best scores in all cognitive domains (Figure 6; S8 Table). In multivariate analysis adjusted by the demographics and clinical parameters that differed between clusters (S9 Table), C3 retained better global, executive, and motor performance compared to C2, and better attention/working memory compared to C1. C1 had better motor performance compared to C2 (S9 Table; model’s p<0.05 for all).

**Figure 6.**
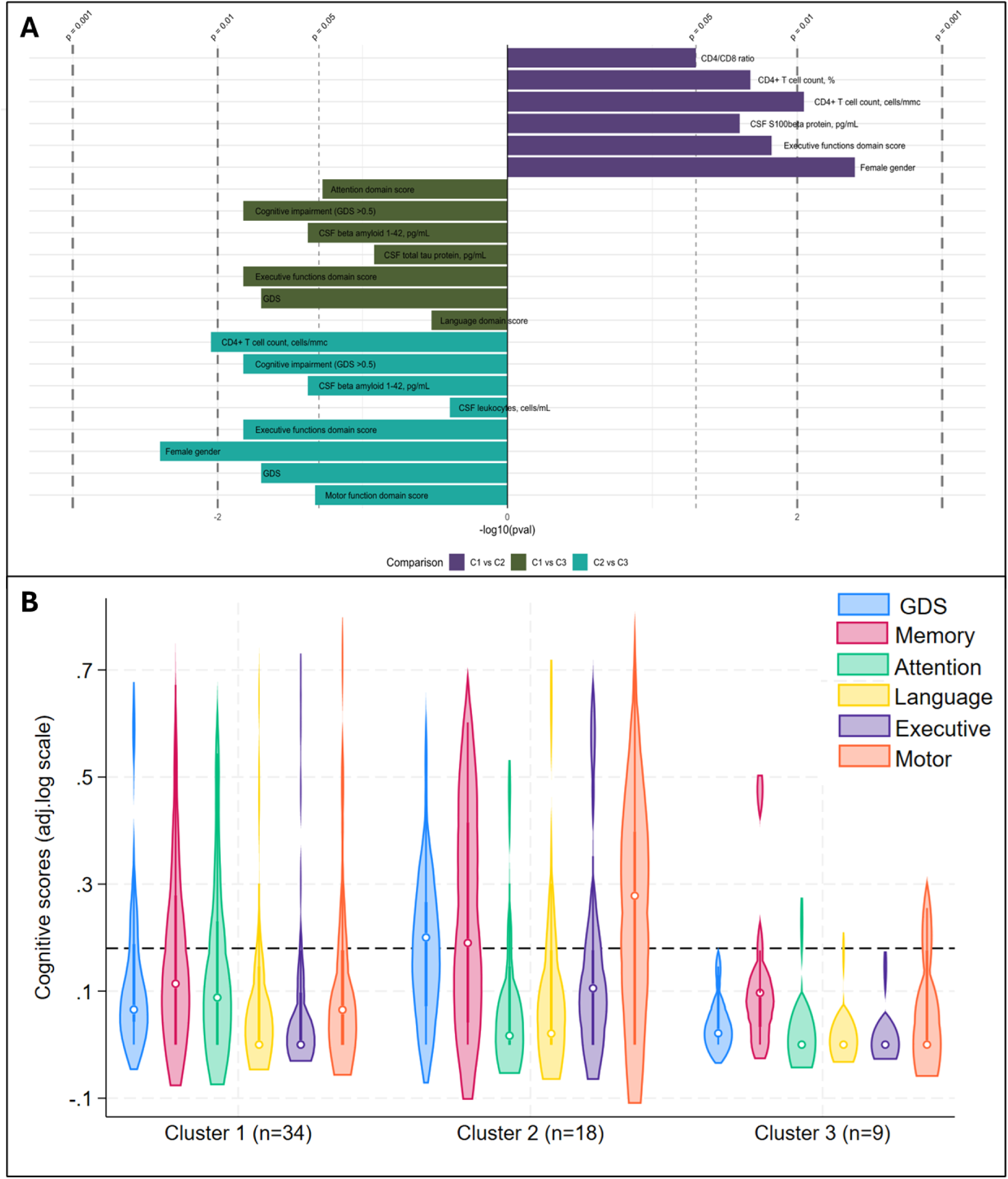
Significant differences between CSF clusters and Cognitive phenotypes of CSF clusters Panel. **A** shows significant differences between clusters: the length of bars represents the size of significance according to p values, while the side of the bar indicates the cluster where the variable is increased/more common (first comparison C2 on the right, second and third comparison C3 on the right). **Panel B** shows violin plots for the Global and Domain Deficit Scores across the clusters; the dotted line cuts at 0.5, threshold to discriminate between impaired (above) and non-impaired cognitive performance (below).

## Discussion

This study provides the first comprehensive description of the CSF virome in PWH. Contrary to our hypothesis, HIV had a weak influence on the virome composition and richness in a population of people with viral suppression and normal CD4 T cell counts. However, only HIV status was associated to the composition of CSF viral communities, while no other demographic or HIV parameter was significantly linked to diversity and composition. We found associations between the cognitive performance and relative abundance of CSF phages, human, and non-human eukaryotic viruses. Neuroinflammation and neurodegeneration may play a role in these relationships as we also found associations between the relative abundance of these viral groups and markers of astrocyte activation, tau protein, and β amyloid.

Approximately half of the CSF samples contained viral sequences. Of these, 93.5% contained PV sequences, and about half hEV or nhEV. Overall, 83.8% of the CSF reads were from viruses that do not infect human cells, primarily phages, with smaller contributions from plant, algae, and fungal viruses. Among the hEV, most belonged to *Herpesviridae*, though we also detected viruses not known to infect cells within or migrating through the CNS. After contig assembly, the number of positive samples and viral diversity decreased, but non-neurotropic viruses still predominated, as 87.5% of contigs were from viruses that primarily infect bacteria, epithelial cells, hepatocytes, or plants.

Participants with CSF phages had no clinical or CSF biochemistry evidence of bacteria in the CNS. Given the virulence of the bacterial hosts (e.g., *Pseudomonas, Cronobacter, Corynebacterium, Burkholderia* spp), the presence of such bacteria in the CSF is unlikely. As discussed in S1 Material, all the identified phages target bacteria that reside in gut, skin, and lung microbiomes. Thus, most of the CSF virome could originate from the periphery.

The length and weight of the contigs (median 2835 bp, ∼1843 kDa) should rule out the free passage of these sequences across the BBB, as hydrophilic molecules >450 Da hardly cross intact BBB [62–64]. The sequences were also too large for pinocytosis, transcytosis, or filtration by the choroid plexus [65,66]. We hypothesize that trafficking cells that have been infected by or have phagocyted these virions/fragments outside the CNS (e.g., macrophages, T cells) largely contributed to the amount of the viral material in the CSF. This interpretation is in line with the well acknowledged Trojan horse mechanism [67] and further evidence, such as peptidoglycan detection in brain lesions infiltrated with blood-derived leukocytes in people suffering from multiple sclerosis [68]. When considering the reads, some sequences had length and weight small enough to potentially cross both intact and injured BBB. Therefore, the presence of some sequences could also be explained by virions/fragments that escape the mucosal barriers and immunity from peripheral niches (e.g., gut, lung, skin) [5,69]. Once in the bloodstream, these can be filtered by the immune system, and small escaping remnants may break through the BBB. This parallel circuit is supported by previous findings, like the detection of small *Helicobacter pylori* components activating astroglia in brain tissues [70]. Lastly, filamentous phages, like some of the *Siphoviridae* detected in our samples, can adhere to epithelial cells via receptors and actively cross barriers through transcytosis and micropinocytosis [5,69,71].

Our findings are consistent with those of the largest study on the CSF virome in people without HIV [20]. Ghose et al. analyzed 20 CSF samples from participants with various conditions (e.g., encephalitis, dementia, schizophrenia, lymphoproliferative cancers), and found that most sequences were from bacteriophages, despite microbiologically proven absence of bacteria [20]. The authors hypothesized that body sites devoid of bacteria, such as CSF, acquire their phageome primarily from the gut. As the CSF virome was not closely related to that found in the stools, they ultimately found less support for this hypothesis [20]. However, our findings, from a larger cohort with less confounding diagnoses, support this hypothesis, considering that the stool microbiome does not accurately represent that in the gut [72,73], and that the composition of the virome of “sterile” body compartments, such as the CSF and bloodstream, could result from the contribution of the gut virome as well as all other viromes of the body. As example, the three most abundant families that we and Ghose [20] detected in the CSF (*Siphoviridae*, *Myoviridae*, *Podoviridae)* are common residents of the human gut, lungs, oral cavity, and skin [2], as all the phages detected (see S1 Material).

An association between the amount of CSF viral sequences and BBB permeability would have provided indirect evidence supporting this model, but neither CSAR nor the age-adjusted classification of BBB injury were associated with the type and amount of viral sequences. In this regard, we could not adjust our analyses by other relevant factors such as the activity of the glymphatic system [74] and local scavengers (e.g., astrocytes and microglia)[75]. Both could contribute to the removal of viral sequences from the CSF. Furthermore, we could not take into account the aforementioned heterogeneity of the possible mechanisms of CNS entry in our analysis. Thus, the conclusion that the CSF virome is not affected by BBB integrity is premature and requires further investigation. This is also why we included CSAR as one of the clustering variables in the post hoc analysis.

A complementary explanation for the CSF phages is that a brain bacteriome does exist [1]. Branton et al. detected sequences of both bacteria and phages in autopsy-derived cerebral white matter of people with and without HIV [22]. The idea of a brain bacteriota was further supported by positive *in situ* staining of peptidoglycan and controlled experiments of transplantations of the CNS tissues into mice, that proved that the viability of the bacteria [22]. The distinction between a “brain microbiota”, referring to living microorganisms, and a “brain microbiome” hypothesis, meant as a pool of genetic fragments that mirror peripheral microbiomes, remains a key question [1]. As we did not perform analyses to distinguish between genetic fragments and virions (e.g., infectivity/viability), our findings are compatible with both the hypotheses, which are non-mutually exclusive. Longitudinal studies on larger samples from blood, CNS, and other body sites, and including different experimental steps (e.g., nuclease protection assays, transmission electron microscopy) can provide insights into the nature of the CSF viral sequences (e.g., virion-to-fragment ratios), their transient fluctuations or stability over time, and the relationship among the viromes of different body compartments.

We found that HIV infection explains a small part of the variation in CSF viral communities. However, we did not identify unique viral species or other metrics that clearly distinguish the virome of PWH from that of CWH. The small representation of taxa limited the power of this analysis. Larger samples are also required to assess whether, despite viral suppression, HIV infection can decrease the diversity of the CSF virome. We observed lower α diversity in PWH compared to CWH, but the difference did not reach statistical significance. In PWH off ART, with AIDS, or low CD4 T cell count, an expansion of phages and other NHV has been described in the blood [76–79]. On the contrary, ART-mediated viral suppression and CD4 restoration can shift gut and blood virome profiles towards that of people without HIV [27,31,78], which can suggest a true lack of association between alpha diversity and HIV status due to the good viroimmunological profile of our study population, and explains the weak effect size of HIV on the CSF virome composition. Direct comparisons of our findings with prior studies are hindered by the differences in the anatomical sites, demographic and viro-immunological characteristics of the study populations, including sexual behaviors [78], and sequencing methods [25,27,30,31,76–78,80,81]. As example, a richer virome has been described in postmortem prefrontal cortex tissues of PWH compared to CWH [25]. The difference with our findings can be ascribed to differences between *in vivo* and postmortem samples, study populations, sequencing pipelines, and to the possibility that the CSF and brain viromes may not be closely related despite anatomical proximity.

Our second hypothesis was that the presence of viral sequences in the CSF contributes to neuroinflammation and affects the brain and mental health of PWH. The etiology behind the HIV-associated neuroinflammation and increased risk of cognitive and mood problems is multifactorial and not fully elucidated [40,82,83], with direct contributions of HIV less important during ART [84–87]. For example, the intrathecal synthesis of immunoglobulins is a common feature of HIV-associated cognitive problems and primarily driven by non-HIV antigens [88–90]. We and others have shown that immune responses against common viruses (e.g., HSV-1, EBV, CMV) contribute to neuroinflammation and cognitive impairment in PWH [53–55]. Evidence in the general population also supports the role of several viral infections in mental and brain health [46–52]. To date, however, the role of the CSF virome in neurocognitive problems has not been investigated. Contrary to our hypothesis, we found no association between the total amount of CSF viral sequences and neuroinflammation, neurodegeneration, cognition, or depressive symptoms. Intrathecal synthesis was also not associated with any characteristics of the virome. As CSF immunoglobulins mostly originate in brain-resident cells and not CSF floating cells [88], this lack of association may reflect weak quantitative and qualitative overlap between the CSF and brain virome. While we measured common pathways of neuroinflammation and neurodegeneration, we may have missed others (e.g., short viral sequences can trigger IFNγ production [91,92], previously linked to neurocognitive issues and neuroinflammation in PWH [93]).

Instead, we found that the RA of hEV, nhEV, and PV was associated with markers of neurodegeneration (βA-42, total tau), astrocyte activation, and cognitive functions. Higher RA of hEV was linked to better global and across-domain cognitive performance as well as to lower βA-42 levels. In contrast, higher RA of nhEV correlated with worse global performance and executive functions, and higher RA of PV with higher βA-42 and S100β levels. The increased activation of astrocytes, represented by increased S100β [94], suggests astrocytic responses to PV. Accordingly, cluster analysis showed greater astrocytosis in participants with NHV-enriched CSF compared to those with virus-devoid CSF. The role of S100β and astrocytes in response to non-human viruses requires further validation considering the phagocytic activity [95,96] and involvement in antiviral innate immune responses [97] of these cells.

Interactions between viruses and βA-42 have been previously described [98–100]. Longitudinal studies should assess the trajectory and outcomes of the relationship herein observed, as βA-42 levels can increase to contain viruses, but when chronically stimulated, they can reduce due to amyloid plaque deposition [98–100]. In our study, only 5 PWH had abnormally low βA-42 levels, as seen in Alzheimer’s dementia; therefore, the association of CSF βA-42 levels with the RA of hEV and PV should be interpreted within a non-pathological range and suggests that PV sequences in the CSF may be a stronger trigger for βA-42 production compared to hEV or nhEV. At post hoc analyses, PWH with hEV-enriched CSF had better cognitive performance and lower βA-42 compared to participants with NHV-enriched or virus-devoid CSF. While this seems opposite to the evidence that chronic hEV infection can contribute to neurodegenerative disorders and neuroinflammation [46–55], the role of hEV has never been investigated alongside other virome components. Using C3 as the reference “expected” CSF virome, the absence of viral sequences in C1 may reflect robust immune responses that clear the CSF thoroughly, but come along with higher inflammation, neuronal injury (the higher total tau), and cognitive disturbance (the worse scores in attention, language, and executive functions). On the contrary, both C2 and C3 had their CSF virome populated by hEV and NHV, but NHV predominance in C2 may indicate impaired CSF clearance or “viral dysbiosis”, for which normative references and contributors are unknown [2,4]. Under the “brain microbiome” hypothesis, NHV should not persist in the CSF, lacking strategies and viable hosts; thus, their presence may result from impaired mechanisms of local clearance (e.g., glymphatic system, phagocytosis of glial cells). Notably, dysfunction of both the glymphatic system and astroglial phagocytosis have been linked to cognitive impairment [95,96,101–103]. Thus, the CSF virome composition could be an epiphenomenon of the true causes of cognitive problems rather than the cause itself. Similarly, C2 had more CSF mononuclear leukocytes than C3. An influx of specific blood leukocytes (e.g., intermediate monocytes) has been linked to larger HIV reservoir in the CNS, neuroinflammation, and cognitive impairment [104,105]. The reason for such influx in some but not all PWH is unknown. The higher RA of NHV in C2 and its worse cognitive performance may be due to enhanced influx of blood leukocytes harboring viral sequences into the CNS, or vice versa the higher RA of NHV may be the driver of the increase leukocyte afflux. Also in this case, the CSF virome could be either the result of other processes that underly cognitive issues or directly contribute to them. Further studies are needed to provide normative references for the human CSF virome and viral dysbiosis, and help disentangling the causality in our associations.

The main limitation of the study is the cross-sectional design that limits causal inferences and directionality in the associations; we cannot conclude whether the difference in CSF viral communities occur during or due to HIV infection, nor whether the CSF virome composition actively contributes to cognitive performance and neuroinflammation or the bystander of concurrent processes responsible of both. Among other limitations, the retrospective nature of the analysis may have affected the detection of genetic material, particularly RNA, due to the long-term preservation of some samples. Although we condensed our analyses to four runs to minimize potential batch effects, we cannot completely rule them out. The virus-like particle enrichment step was implemented to reduce genetic background interference and enhance viral recovery in fluids expected to be poor in viral sequences, as previously suggested [20,25,106]. However, this step could affect viral diversity by missing less abundant, latently infecting, integrated viruses, and larger viruses. Our sequencing depth was unable to detect low-level sequences from HIV, EBV, and JCV, identified via RT-PCR, as expected for specimens with <100 cp/mL [59–61]. Thus, the richness and composition of the virome is likely underestimated. All samples were positive for viral reads prior to applying confidence thresholds (≥two unique reads and Kraken confidence score >0.5). Although this approach strengthens confidence in the taxonomic attribution, it may lead to excluding sequences due to their scarcity, missing matches in the reference database, or uncertain attribution. While grouping CSF viruses by target hosts fits with our hypothesis model, it may oversimplify the heterogeneity of effects and characteristics within each group. Lastly, CWH had no neurocognitive assessment limiting the possibility to include this group in all the analyses.

Among the strengths, the unbiased approach of shotgun sequencing and the post hoc clustering analyses, the comprehensive assessment of cognition, mood, and biomarkers in the largest set of CSF samples used for shotgun viral metagenomics, the comparison of an assembly-free vs assembly-based method, the ecological validity of our study population in representing modern PWH, and the inclusion of internal negative control specimens and CWH. The use of blank controls may have resulted in the exclusion of some taxa truly present, but it ensured the highest confidence in the absence of contamination during the experiments. The risk of pre-analytical contamination was also minimized by sterilizing the skin at the LP site, using the last collected CSF sample (∼7 tubes) for sequencing, and confirming the absence of red blood cells in all samples. Lastly, the use of specimens from living participants, rather than postmortem samples, avoids inevitable contaminations occurring after death, and provides insights on *in vivo* phenomena not confounded by postmortem processes.

In conclusion, this study offers the first comprehensive description of the CSF virome of PWH on suppressive ART and valid CD4 T cell count. HIV infection exerted only a modest influence on the CSF virome, despite being the unique factor associated with the composition of viral communities. Most CSF sequences were from viruses that have no recognized host within the CNS, suggesting a peripheral source for most of these sequences. Half belonged to bacteriophages commonly found in other human viromes, followed by equal proportions of human and non-human eukaryotic viruses. The relative abundance of these viral groups was associated with cognition in PWH, with higher abundance of non-human eukaryotic viruses associated with poorer cognitive performance. Distinct viral landscapes were also linked to diverse signatures of neuroinflammation and neurodegeneration, with higher abundance of phages associated with astrocytosis and of human eukaryotic viruses with lower amyloid β. Further investigation is warranted to understand the mechanisms driving these relationships, identify the origin of the CSF virome, and explore its role in brain and mental health.

## Materials and Methods

### 1. Study design and population

We performed a pilot cross-sectional study in 85 CSF samples from 74 adult PWH and 11 adults CWH to describe the presence and composition of the CSF virome and to investigate its relationships HIV infection, neuroinflammation, depressive symptoms, and cognitive performance.

CSF samples from PWH were collected and stored between January 2010 and September 2019 at the Unit of Infectious Diseases, Amedeo di Savoia hospital, Torino (Italy) and retrospectively analyzed for CSF virome through shotgun metagenomics. From the CSF library, CSF samples of ≥1 mL were selected based on the following: A) Mandatory inclusion criteria: age ≥18 years, plasma HIV-RNA<200 cp/mL, being on standard three-drug NRTI-based therapy (2 NRTIs + either one protease inhibitor, integrase strand-transfer inhibitor, or non-nucleoside reverse transcriptase inhibitor) since at least 12 months, and signed consent for the storage and use of samples and data for future analyses at the time of lumbar puncture; B) Preferential inclusion criteria: biomarkers of interest already measured in CSF collected at the same time of the sample used for the current analysis, neurocognitive and mood assessment at the time of CSF collection (±1 month), balanced female to male ratio; C) Exclusion criteria: diagnosis of major confounding factors for cognitive evaluation and CSF biomarkers prior or at CSF collection (e.g., CNS disorders or infections, language barriers, cerebrovascular events, head trauma, neuropathy), and substance/alcohol use disorders in the previous year.

A control group of 11 CWH was enrolled between March and June 2023. HIV-1/2 infection was ruled out at the time of CSF collection through fourth-generation commercial serology assay. Participation was prospectively offered by the Unit of Emergency Medicine, Reanimation and Anesthesia at Maria Vittoria hospital, Torino (Italy), to adult individuals who had to undergo spinal anesthesia for surgical indications, with negative medical history for immunological and CNS disorders (e.g., acquired or congenital immune deficits, cancer, immuno-rheumatological diseases, neurodegenerative disorders) and for active infections. All CWH signed an informed consent for the storage and analyses of CSF collected during spinal anesthesia. Spinal anesthesia was required for orthopedic (3 femur fractures or coxarthrosis, 2 meniscus injury), urology (1 transurethral prostate resection, 1 varicocelectomy), and abdominal surgery (2 hernioplasty, 1 intervention for hemorrhoids, 1 polyps’ removal).

The research was performed in accordance with the Declaration of Helsinki and has been approved by the Comitato Etico Interaziendale A.O.U. Città della Salute e della Scienza di Torino, A.O. Ordine Mauriziano di Torino, A.S.L. Città di Torino (protocol n.285/2022).

### 2. Clinical assessment and CSF biomarkers

Standardized assessments and physical examination were performed at the time of CSF collection to record data on demographics, clinical, HIV disease characteristics, and biomarkers.

Stored CSF samples were frozen (-80°C) within 1 hour from the lumbar puncture, while immediately after collection paired CSF was analyzed for: standard biochemistry (cells, glucose, and protein); markers of neurodegeneration (total tau, 181-phosphorylated tau [ptau], and fragment 1–42 of β amyloid [Aβ-42]) through commercial immunoassays (Innogenetics, Ghent, Belgium, EU); markers of neuro-inflammation (neopterin and S100β protein) through commercial immunoassays (DRG Diagnostics, Marnurg, Germany; DIAMETRA Srl, Spello, Italy); BBB integrity through the CSF-to-serum albumin ratio (CSAR, CSF albumin mg/L:serum albumin mg/L); and intrathecal synthesis by IgG index (CSF IgG:serum IgG:CSAR) according to age-adjusted Reibergrams [133].

Cut-offs for normality of the CSF biomarkers were based on standardized routine cut-offs (e.g., CSF cells>5/mL; CSAR <6.5 in subjects aged <40 years and <8 in older subjects) and manufacturer’s cut-offs for the general population, and previously used in PWH [86,107,108]: tau >300 pg/mL (age ≤50 years), >450 pg/mL (age 51–70 years), >500 pg/mL (age > 70 years); ptau >61 pg/mL; Aβ42 <500 pg/mL; neopterin >1.5 ng/mL; S100β >380 pg/mL.

HIV RNA was quantified in plasma and CSF by reverse-transcription polymerase chain reaction (Roche Amplicor, lower limit of quantitation 20 copies/mL). Results of PCR for EBV DNA, CMV DNA, and JCV DNA (in-house standardized real-time PCR) were available in the CSF of a subsample of PWH (S2 Table).

### 3. Mood and Cognitive Assessment

A subgroup of PWH (n=61) underwent a comprehensive battery of cognitive tests within ±1 month from CSF collection. Raw scores were converted to demographically corrected T scores by referencing to published normative standards which correct for age, education, and sex. Five cognitive domains were assessed by 13 tests (S8 Table): attention/working memory, abstraction/executive functions, short- and long-term memory (auditory and visual), verbal fluency/language, and motor functions. Individual domain scores were combined to calculate the Global Deficit Score (GDS), as previously described [109]; a GDS ≥0.5 indicated neurocognitive impairment [109].

Depressive mood was assessed in a subgroup of PWH (n=66) through the Beck Depression Inventory II (BDI-II) [110]. Cut-off for the severity of depressive symptoms have been validated in the general population, but are commonly used in PWH [111,112]: a total score ≤13 is considered absent/subthreshold depression, 14–19 mild, 20–28 moderate, and 29–63 severe depressive symptoms.

### 4. Shotgun metagenomics

#### 4.1 CSF samples preparation and nucleic acid extraction

Prokaryotic and eukaryotic viruses in CSF were first enriched by isolation and purification of virus-like particles. This purification was performed through a filtration process that takes advantage of the smaller dimension of viruses with respect to eukaryotic and prokaryotic cells. CSF samples were initially filtered with 0.45 µm polyethersulfone filters (Merck millipore) to retain most of the bacteria and cellular contaminants and then with 0.22 µm polyethersulfone filters (Merck millipore) capable of enriching viruses.

The resulting filtrate (∼200-400 µl of sample) was used for nucleic acid extraction using the QIAamp MinElute Virus Spin Kit (Qiagen, Hilden, Germany), according to the manufacturer’s instructions. Both the concentration and the quality of the extracted nucleic acids were assessed by Nanodrop.

#### 4.2 Amplification and sequencing

One hundred ng input of purified nucleic acids was used for genome amplification and sequencing following a modification of the QIAseq® Single Cell RNA Library Kits with Unique Dual Indexes protocol. Briefly, a first reverse transcription, amplification and cDNA production were performed from total viral nucleic acids (RNA plus DNA) to obtain a pool of cDNA. The cDNA obtained from the previous step was first subjected to enzymatic fragmentation and then used for libraries preparation by using the QIAseq Single Cell RNA Amplified cDNA. The produced libraries were purified with QIAseq Beads, were validated using the High Sensitivity D1000 ScreenTape system on a Bioanalyzer (Agilent Technologies), and then quantified using a High Sensitivity Double Stranded DNA kit on a Qubit Fluorometer (Thermo Fisher Scientific). Normalized indexed DNA libraries were then loaded onto an Illumina High Output Flow cell cartridge v.2.5, and sequenced using the NextSeq 550 instrument (Illumina, San Diego, CA, USA) with 2×150-bp paired-end reads.

Three different runs for the CSF samples from PWH, and a fourth run for the CSF samples from CWH were performed. In each run, the blank samples were processed in parallel with CSF samples to document contamination background.

#### 4.3 Virome characterization

Demultiplexed raw reads were trimmed for adapter and quality (Phred score >28) and deduplicated using Fastp (v0.23.2) [113]. The final read quality was assessed by FastQC (v0.11.9) and MultiQC (v1.12) [114,115]. All reads mapping to human genome (GRCh38) were removed from all the samples using bbsplit (BBTools). The remaining clean reads were transformed back to paired end reads using reformat (BBTools). Taxonomy was assigned using Kraken 2 (v2.1.2) [116] and visualized by Krona (v2.8.1) [117], while Bracken (v2.7) [118] was used to estimate family abundance.

The sequencing results underwent two parallel methods of taxonomic assignment: based on reads and based on contigs reconstruction. Reads classified as viral (taxonID 10239) were retrieved using the ‘extract_kraken_reads.py’ script of the KrakenTools suite v1.2. Taxonomy assignment was obtained by supporting each viral taxon with at least two unique reads and a Kraken confidence score >0.50, as suggested [119,120]. Species observed from the blanks were removed from all samples (for details see S9 Table).

For contigs reconstruction, reads retained after the previous steps were *de-novo* assembled using metaSPAdes (v3.14.1) [121] in paired-read mode with default settings. The obtained contigs were filtered for length ≥300bp, as suggested [122], using reformat from BBTools, and then queried against a nucleotide viral sequences collection retrieved from the National Center for Biotechnology Information (NCBI) Reference Sequence Database (RefSeq, 8416 sequences) using Nucleotide BLAST (blastn) with an e-value of at least 1×10^-10^ and a percentage identity of at least 70%. Species observed by the contigs assembled from the blanks (without length filtering) were removed from all samples (for details see S9 Table).

Eukaryotic and non-eukaryotic viral species and their genome coverage were visualized through a heatmap constructed using the ggplot2 (v3.3.6) and pheatmap (v1.0.12) R packages [123]. The filtered Bracken report output files were used to generate a BIOM-format table using the kraken-biom python package (v1.0.1). The biom-format package (v2.1.12) [124] was then used to add sample metadata to the BIOM table, which was then loaded into R studio. The sequences used in this work have been deposited in GenBank (released date 01/31/25; BioProject ID PRJNA1176451).

### 5. Statistical Analyses

Data were presented as median (interquartile range, IQR), mean (standard deviation, SD), and number (proportion, %), as appropriate. Non-normally distributed variables (e.g., HIV RNA, biomarker levels) were log10-transformed to reduce skewness. Missing data accounted for a maximum of 5.4% of the observations across 11 variables for PWH (reported in S10 Table) and were addressed by excluding the affected participants in the corresponding analyses.

We characterized alpha diversity using the number of observed taxa (richness of samples), the Simpson index (which is more sensitive to the dominance of species), and the Shannon index (which is sensitive to both richness of species and evenness of their distribution). We characterized the beta diversity using Bray Curtis dissimilarity (occurrence and abundance of taxa) and Jaccard distance (presence/absence of taxa), tested the significance through Permutation Based Analysis of Variance (PERMANOVA) with Adonis, and visualized the output using Principal Coordinate Analysis (PCoA). The following characteristics of CSF virome were also investigated and referred as viral metrics: total number of viral reads per CSF sample and relative abundance of viral categories (number of reads of each category divided by total number of viral reads per sample). HEV, nhEV, and PV were used as viral categories; we did not proceed to assess single viral families due to the limited number of positive samples for each family.

Kruskal-Wallis rank sum test, Mann-Whitney U test, Fisher’s exact test, Chi-squared test, and Pearson and Spearman’s rank correlations were used based on data distribution and variables type. Hierarchical clustering was performed to identify distinct groups within the dataset; the distance matrix was computed using Euclidean distance, as commonly suggested for continuous data [125]. Prior to clustering, input data were log-transformed to stabilize variance and improve comparability across variables. Hierarchical clustering was conducted using Ward’s minimum variance method (Ward.D2), which minimizes the total within-cluster variance at each step and optimizes cluster formation [126]. The resulting dendrogram was visualized to assess cluster structure, and the final number of clusters was selected based on visual inspection [127]. Generalized linear models were used to confirm the associations between CSF clusters and cognitive performance after considering other factors that significantly differed between the clusters.

Negative results after sequencing can be biological (meaningful information) or non-biological zeros (e.g., due to limits in sequencing depth, degradation of genetic material) [128]. All the CSF samples were included in the analyses investigating quantitative relationships between viral metrics and other variables of interest (e.g., correlations with cognitive scores and biomarkers), considering as closely related real biological zeros and samples containing levels of viral genetic material below the limit of detection of sequencing. However, taking into consideration the wide range of conservation time of the specimens and the differential stability of DNA versus RNA, among possible factors contributing to negative measurements, we also performed sensitivity analyses in virome-positive samples only, to assess how the observed associations changed with and without including the negative samples. Analyses were performed through RStudio v4.3.3 (R Core Team 2024, Boston, MA, US) and STATA v18 (StataCorp, College Station, TX, US).

## Acknowledgments

We are grateful to the participants, to those who provided technical assistance and collaborators, and to the funding agency (Gilead Sciences Srl, Italy).

## Supporting information captions

**S1 Table.** Demographic and HIV-related characteristics

**S2 Table.** Viruses detected in the blank samples and removed from the study samples

**S3 Table.** Comparison of CSF viruses detected by PCR and Sequencing

**S4 Table.** Clinical characteristics of the 8 participants with CSF positive for eukaryotic viruses after contigs reconstruction

**S5 Table.** Clinical characteristics of the 5 participants with CSF positive for prokaryotic viruses after contigs reconstruction

**S1 Material.** Case-series description of the 12 participants positive for CSF viruses after contigs reconstruction

**S6 Table.** Results of CSF biomarkers and neurocognitive assessment

**S7 Table.** Correlations between relative abundance of CSF viral categories and neurocognitive measures

**S8 Table.** Comparisons of Demographic, HIV-related parameters, Neuroinflammation and Neurocognitive measures between Clusters

**S9 Table.** Multivariate analysis for neurocognitive metrics

**S10 Table.** Battery of cognitive tests used for neurocognitive assessment

**S11 Table.** Missing data

**S12 Anonymized Study Dataset**

## Data Availability

The sequences used in this work have been deposited in GenBank (released date 01/31/25; BioProject ID PRJNA1176451). All other relevant data and metadata supporting the findings of this study are available online as online Supporting Information 12 after removal of direct and indirect identifiers.

## References

1. Link CD. Is There a Brain Microbiome? Neurosci Insights. 2021;16: 26331055211018709. doi:10.1177/26331055211018709

2. Liang G, Bushman FD. The human virome: assembly, composition and host interactions. Nat Rev Microbiol. 2021;19: 514–527. doi:10.1038/s41579-021-00536-5

3. Moustafa A, Xie C, Kirkness E, Biggs W, Wong E, Turpaz Y, et al. The blood DNA virome in 8,000 humans. PLoS Pathog. 2017;13: e1006292. doi:10.1371/journal.ppat.1006292

4. Rascovan N, Duraisamy R, Desnues C. Metagenomics and the Human Virome in Asymptomatic Individuals. Annu Rev Microbiol. 2016;70: 125–141. doi:10.1146/annurev-micro-102215-095431

5. Cao Z, Sugimura N, Burgermeister E, Ebert MP, Zuo T, Lan P. The gut virome: A new microbiome component in health and disease. eBioMedicine. 2022;81. doi:10.1016/j.ebiom.2022.104113

6. Bai G-H, Lin S-C, Hsu Y-H, Chen S-Y. The Human Virome: Viral Metagenomics, Relations with Human Diseases, and Therapeutic Applications. Viruses. 2022;14: 278. doi:10.3390/v14020278

7. Popescu M, Van Belleghem JD, Khosravi A, Bollyky PL. Bacteriophages and the Immune System. Annu Rev Virol. 2021;8: 415–435. doi:10.1146/annurev-virology-091919-074551

8. Castells-Nobau A, Mayneris-Perxachs J, Fernández-Real JM. Unlocking the mind-gut connection: Impact of human microbiome on cognition. Cell Host Microbe. 2024;32: 1248–1263. doi:10.1016/j.chom.2024.07.019

9. Mayneris-Perxachs J, Castells-Nobau A, Arnoriaga-Rodríguez M, Garre-Olmo J, Puig J, Ramos R, et al. Caudovirales bacteriophages are associated with improved executive function and memory in flies, mice, and humans. Cell Host Microbe. 2022;30: 340–356.e8. doi:10.1016/j.chom.2022.01.013

10. Shkoporov AN, Turkington CJ, Hill C. Mutualistic interplay between bacteriophages and bacteria in the human gut. Nat Rev Microbiol. 2022;20: 737–749. doi:10.1038/s41579-022-00755-4

11. Neil JA, Cadwell K. The Intestinal Virome and Immunity. J Immunol. 2018;201: 1615–1624. doi:10.4049/jimmunol.1800631

12. Kviatcovsky D, Valdés-Mas R, Federici S, Elinav E. Phage therapy in noncommunicable diseases. Science. 2023;382: 266–267. doi:10.1126/science.adh2718

13. Wahida A, Tang F, Barr JJ. Rethinking phage-bacteria-eukaryotic relationships and their influence on human health. Cell Host & Microbe. 2021;29: 681–688. doi:10.1016/j.chom.2021.02.007

14. Ritz NL, Draper LA, Bastiaanssen TFS, Turkington CJR, Peterson VL, van de Wouw M, et al. The gut virome is associated with stress-induced changes in behaviour and immune responses in mice. Nat Microbiol. 2024;9: 359–376. doi:10.1038/s41564-023-01564-y

15. Yolken RH, Jones-Brando L, Dunigan DD, Kannan G, Dickerson F, Severance E, et al. Chlorovirus ATCV-1 is part of the human oropharyngeal virome and is associated with changes in cognitive functions in humans and mice. Proc Natl Acad Sci U S A. 2014;111: 16106–16111. doi:10.1073/pnas.1418895111

16. Levine KS, Leonard HL, Blauwendraat C, Iwaki H, Johnson N, Bandres-Ciga S, et al. Virus exposure and neurodegenerative disease risk across national biobanks. Neuron. 2023;111: 1086–1093.e2. doi:10.1016/j.neuron.2022.12.029

17. Bjornevik K, Münz C, Cohen JI, Ascherio A. Epstein–Barr virus as a leading cause of multiple sclerosis: mechanisms and implications. Nat Rev Neurol. 2023;19: 160–171. doi:10.1038/s41582-023-00775-5

18. de Haan L, Sutterland AL, Schotborgh JV, Schirmbeck F, de Haan L. Association of Toxoplasma gondii Seropositivity With Cognitive Function in Healthy People. JAMA Psychiatry. 2021;78: 1–10. doi:10.1001/jamapsychiatry.2021.1590

19. Davis AP, Maxwell CL, Mendoza H, Crooks A, Dunaway SB, Storey S, et al. Cognitive impairment in syphilis: Does treatment based on cerebrospinal fluid analysis improve outcome? PLoS One. 2021;16: e0254518. doi:10.1371/journal.pone.0254518

20. Ghose C, Ly M, Schwanemann LK, Shin JH, Atab K, Barr JJ, et al. The Virome of Cerebrospinal Fluid: Viruses Where We Once Thought There Were None. Front Microbiol. 2019;10: 2061. doi:10.3389/fmicb.2019.02061

21. Blanco-Picazo P, Fernández-Orth D, Brown-Jaque M, Miró E, Espinal P, Rodríguez-Rubio L, et al. Unravelling the consequences of the bacteriophages in human samples. Sci Rep. 2020;10: 6737. doi:10.1038/s41598-020-63432-7

22. Branton WG, Ellestad KK, Maingat F, Wheatley BM, Rud E, Warren RL, et al. Brain microbial populations in HIV/AIDS: α-proteobacteria predominate independent of host immune status. PLoS One. 2013;8: e54673. doi:10.1371/journal.pone.0054673

23. Ghorbani M. Unveiling the Human Brain Virome in Brodmann Area 46: Novel Insights Into Dysbiosis and Its Association With Schizophrenia. Schizophrenia Bulletin Open. 2023;4: sgad029. doi:10.1093/schizbullopen/sgad029

24. Pyöriä L, Pratas D, Toppinen M, Hedman K, Sajantila A, Perdomo MF. Unmasking the tissue-resident eukaryotic DNA virome in humans. Nucleic Acids Res. 2023;51: 3223–3239. doi:10.1093/nar/gkad199

25. Dang X, Hanson BA, Orban ZS, Jimenez M, Suchy S, Koralnik IJ. Characterization of the brain virome in human immunodeficiency virus infection and substance use disorder. PLoS One. 2024;19: e0299891. doi:10.1371/journal.pone.0299891

26. Ortiz AM, Brenchley JM. Untangling the role of the microbiome across the stages of HIV disease. Curr Opin HIV AIDS. 2024;19: 221–227. doi:10.1097/COH.0000000000000870

27. Villoslada-Blanco P, Pérez-Matute P, Íñiguez M, Recio-Fernández E, Jansen D, De Coninck L, et al. Impact of HIV infection and integrase strand transfer inhibitors-based treatment on the gut virome. Sci Rep. 2022;12: 21658. doi:10.1038/s41598-022-25979-5

28. Lippincott RA, O’Connor J, Neff CP, Lozupone C, Palmer BE. Deciphering HIV-associated inflammation: microbiome’s influence and experimental insights. Curr Opin HIV AIDS. 2024;19: 228–233. doi:10.1097/COH.0000000000000866

29. Ishizaka A, Koga M, Mizutani T, Parbie PK, Prawisuda D, Yusa N, et al. Unique Gut Microbiome in HIV Patients on Antiretroviral Therapy (ART) Suggests Association with Chronic Inflammation. Microbiology Spectrum. 2021;9: 10.1128/spectrum.00708-21.

30. Monaco CL, Gootenberg DB, Zhao G, Handley SA, Ghebremichael MS, Lim ES, et al. Altered Virome and Bacterial Microbiome in Human Immunodeficiency Virus-Associated Acquired Immunodeficiency Syndrome. Cell Host Microbe. 2016;19: 311–322. doi:10.1016/j.chom.2016.02.011

31. Monaco CL. HIV, AIDS, and the virome: Gut reactions. Cell Host Microbe. 2022;30: 466–470. doi:10.1016/j.chom.2022.03.010

32. Fulcher JA, Li F, Tobin NH, Zabih S, Elliott J, Clark JL, et al. Gut dysbiosis and inflammatory blood markers precede HIV with limited changes after early seroconversion. eBioMedicine. 2022;84. doi:10.1016/j.ebiom.2022.104286

33. Bai X, Nielsen SD, Kunisaki KM, Trøseid M. Pulmonary comorbidities in people with HIV-the microbiome connection. Curr Opin HIV AIDS. 2024;19: 246–252. doi:10.1097/COH.0000000000000871

34. Bulnes R, Utay NS. Therapeutic microbiome modulation: new frontiers in HIV treatment. Curr Opin HIV AIDS. 2024;19: 268–275. doi:10.1097/COH.0000000000000864

35. Lam JO, Hou CE, Hojilla JC, Anderson AN, Gilsanz P, Alexeeff SE, et al. Comparison of dementia risk after age 50 between individuals with and without HIV infection. AIDS. 2021;35: 821–828. doi:10.1097/QAD.0000000000002806

36. Hyle EP, Wattananimitgul N, Mukerji SS, Foote JHA, Reddy KP, Thielking A, et al. Age-associated dementia among older people aging with HIV in the United States: a modeling study. AIDS. 2024;38: 1186–1197. doi:10.1097/QAD.0000000000003862

37. Wang Y, Liu M, Lu Q, Farrell M, Lappin JM, Shi J, et al. Global prevalence and burden of HIV-associated neurocognitive disorder: A meta-analysis. Neurology. 2020;95: e2610–e2621. doi:10.1212/WNL.0000000000010752

38. Trunfio M, Vai D, Montrucchio C, Alcantarini C, Livelli A, Tettoni MC, et al. Diagnostic accuracy of new and old cognitive screening tools for HIV-associated neurocognitive disorders. HIV Med. 2018. doi:10.1111/hiv.12622

39. Du X, Zhang Q, Hao J, Gong X, Liu J, Chen J. Global trends in depression among patients living with HIV: A bibliometric analysis. Front Psychol. 2023;14: 1125300. doi:10.3389/fpsyg.2023.1125300

40. Mudra Rakshasa-Loots A, Whalley HC, Vera JH, Cox SR. Neuroinflammation in HIV-associated depression: evidence and future perspectives. Mol Psychiatry. 2022;27: 3619–3632. doi:10.1038/s41380-022-01619-2

41. Wojna V. Neuroinflammation and HIV-Associated Cognitive Impairment. Neurology. 2022;99: 499–500. doi:10.1212/WNL.0000000000201095

42. Carrico AW, Cherenack EM, Rubin LH, McIntosh R, Ghanooni D, Chavez JV, et al. Through the Looking-Glass: Psychoneuroimmunology and the Microbiome-Gut-Brain Axis in the Modern Antiretroviral Therapy Era. Psychosom Med. 2022;84: 984–994. doi:10.1097/PSY.0000000000001133

43. Hua S, Peters BA, Lee S, Fitzgerald K, Wang Z, Sollecito CC, et al. Gut Microbiota and Cognitive Function Among Women Living with HIV. J Alzheimers Dis. 2023;95: 1147–1161. doi:10.3233/JAD-230117

44. Rich S, Klann E, Bryant V, Richards V, Wijayabahu A, Bryant K, et al. A review of potential microbiome-gut-brain axis mediated neurocognitive conditions in persons living with HIV. Brain Behav Immun Health. 2020;9: 100168. doi:10.1016/j.bbih.2020.100168

45. Chen X, Wei J, Zhang Y, Zhang Y, Zhang T. Crosstalk between gut microbiome and neuroinflammation in pathogenesis of HIV-associated neurocognitive disorder. J Neurol Sci. 2024;457: 122889. doi:10.1016/j.jns.2024.122889

46. Balin BJ, Hudson AP. Herpes viruses and Alzheimer’s disease: new evidence in the debate. The Lancet Neurology. 2018;17: 839–841. doi:10.1016/S1474-4422(18)30316-8

47. Khalesi Z, Tamrchi V, Razizadeh MH, Letafati A, Moradi P, Habibi A, et al. Association between human herpesviruses and multiple sclerosis: A systematic review and meta-analysis. Microb Pathog. 2023;177: 106031. doi:10.1016/j.micpath.2023.106031

48. Readhead B, Haure-Mirande J-V, Funk CC, Richards MA, Shannon P, Haroutunian V, et al. Multiscale Analysis of Independent Alzheimer’s Cohorts Finds Disruption of Molecular, Genetic, and Clinical Networks by Human Herpesvirus. Neuron. 2018;99: 64–82.e7. doi:10.1016/j.neuron.2018.05.023

49. Ma X, Liao Z, Tan H, Wang K, Feng C, Xing P, et al. The association between cytomegalovirus infection and neurodegenerative diseases: a prospective cohort using UK Biobank data. eClinicalMedicine. 2024;74. doi:10.1016/j.eclinm.2024.102757

50. Bjornevik K, Cortese M, Healy BC, Kuhle J, Mina MJ, Leng Y, et al. Longitudinal analysis reveals high prevalence of Epstein-Barr virus associated with multiple sclerosis. Science. 2022;375: 296–301. doi:10.1126/science.abj8222

51. Grut V, Biström M, Salzer J, Stridh P, Jons D, Gustafsson R, et al. Human herpesvirus 6A and axonal injury before the clinical onset of multiple sclerosis. Brain. 2024;147: 177–185. doi:10.1093/brain/awad374

52. Hakami MA, Khan FR, Abdulaziz O, Alshaghdali K, Hazazi A, Aleissi AF, et al. Varicella-zoster virus-related neurological complications: From infection to immunomodulatory therapies. Rev Med Virol. 2024;34: e2554. doi:10.1002/rmv.2554

53. Trunfio M, Sacchi A, Vai D, Pittaluga F, Croce M, Cavallo R, et al. Intrathecal production of anti-EBV Viral Capsid Antigen IgG is associated with Neurocognition and tau pathology in people with HIV. AIDS. 2023. doi:10.1097/QAD.0000000000003775

54. Trunfio M, Di Girolamo L, Ponzetta L, Russo M, Burdino E, Imperiale D, et al. Seropositivity and reactivations of HSV-1, but not of HSV-2 nor VZV, associate with altered blood–brain barrier, beta amyloid, and tau proteins in people living with HIV. J Neurovirol. 2022 [cited 9 Nov 2022]. doi:10.1007/s13365-022-01105-z

55. Letendre S, Bharti A, Perez-Valero I, Hanson B, Franklin D, Woods SP, et al. Higher Anti-Cytomegalovirus Immunoglobulin G Concentrations Are Associated With Worse Neurocognitive Performance During Suppressive Antiretroviral Therapy. Clin Infect Dis. 2018;67: 770–777. doi:10.1093/cid/ciy170

56. Cai C, Tang Y-D, Xu G, Zheng C. The crosstalk between viral RNA- and DNA-sensing mechanisms. Cell Mol Life Sci. 2021;78: 7427–7434. doi:10.1007/s00018-021-04001-7

57. Champagne-Jorgensen K, Luong T, Darby T, Roach DR. Immunogenicity of bacteriophages. Trends in Microbiology. 2023;31: 1058–1071. doi:10.1016/j.tim.2023.04.008

58. Ayling M, Clark MD, Leggett RM. New approaches for metagenome assembly with short reads. Brief Bioinform. 2020;21: 584–594. doi:10.1093/bib/bbz020

59. Miller S, Naccache SN, Samayoa E, Messacar K, Arevalo S, Federman S, et al. Laboratory validation of a clinical metagenomic sequencing assay for pathogen detection in cerebrospinal fluid. Genome Res. 2019;29: 831–842. doi:10.1101/gr.238170.118

60. Bukowska-Ośko I, Perlejewski K, Nakamura S, Motooka D, Stokowy T, Kosińska J, et al. Sensitivity of Next-Generation Sequencing Metagenomic Analysis for Detection of RNA and DNA Viruses in Cerebrospinal Fluid: The Confounding Effect of Background Contamination. Adv Exp Med Biol. 2016 [cited 8 Jan 2024]. doi:10.1007/5584_2016_42

61. Schlaberg R, Chiu CY, Miller S, Procop GW, Weinstock G, Professional Practice Committee and Committee on Laboratory Practices of the American Society for Microbiology, et al. Validation of Metagenomic Next-Generation Sequencing Tests for Universal Pathogen Detection. Arch Pathol Lab Med. 2017;141: 776–786. doi:10.5858/arpa.2016-0539-RA

62. Pardridge WM. A Historical Review of Brain Drug Delivery. Pharmaceutics. 2022;14: 1283. doi:10.3390/pharmaceutics14061283

63. Sun Y, Zabihi M, Li Q, Li X, Kim BJ, Ubogu EE, et al. Drug Permeability: From the Blood-Brain Barrier to the Peripheral Nerve Barriers. Adv Ther (Weinh). 2023;6: 2200150. doi:10.1002/adtp.202200150

64. Meng F, Xi Y, Huang J, Ayers PW. A curated diverse molecular database of blood-brain barrier permeability with chemical descriptors. Sci Data. 2021;8: 289. doi:10.1038/s41597-021-01069-5

65. Lalatsa A, Butt AM. Chapter 3 -Physiology of the Blood–Brain Barrier and Mechanisms of Transport Across the BBB. In: Kesharwani P, Gupta U, editors. Nanotechnology-Based Targeted Drug Delivery Systems for Brain Tumors. Academic Press; 2018. pp. 49–74. doi:10.1016/B978-0-12-812218-1.00003-8

66. Wu D, Chen Q, Chen X, Han F, Chen Z, Wang Y. The blood–brain barrier: Structure, regulation and drug delivery. Sig Transduct Target Ther. 2023;8: 1–27. doi:10.1038/s41392-023-01481-w

67. Santiago-Tirado FH, Doering TL. False friends: Phagocytes as Trojan horses in microbial brain infections. PLoS Pathog. 2017;13: e1006680. doi:10.1371/journal.ppat.1006680

68. Schrijver IA, van Meurs M, Melief MJ, Wim Ang C, Buljevac D, Ravid R, et al. Bacterial peptidoglycan and immune reactivity in the central nervous system in multiple sclerosis. Brain. 2001;124: 1544–1554. doi:10.1093/brain/124.8.1544

69. Tiamani K, Luo S, Schulz S, Xue J, Costa R, Khan Mirzaei M, et al. The role of virome in the gastrointestinal tract and beyond. FEMS Microbiol Rev. 2022;46: fuac027. doi:10.1093/femsre/fuac027

70. Xie J, Cools L, Van Imschoot G, Van Wonterghem E, Pauwels MJ, Vlaeminck I, et al. Helicobacter pylori-derived outer membrane vesicles contribute to Alzheimer’s disease pathogenesis via C3-C3aR signalling. J Extracell Vesicles. 2023;12: e12306. doi:10.1002/jev2.12306

71. Nguyen S, Baker K, Padman BS, Patwa R, Dunstan RA, Weston TA, et al. Bacteriophage Transcytosis Provides a Mechanism To Cross Epithelial Cell Layers. mBio. 2017;8: 10.1128/mbio.01874-17.

72. Yan A, Butcher J, Schramm L, Mack DR, Stintzi A. Multiomic spatial analysis reveals a distinct mucosa-associated virome. Gut Microbes. 2023;15: 2177488. doi:10.1080/19490976.2023.2177488

73. Ahn J-S, Lkhagva E, Jung S, Kim H-J, Chung H-J, Hong S-T. Fecal Microbiome Does Not Represent Whole Gut Microbiome. Cellular Microbiology. 2023;2023: 6868417. doi:10.1155/2023/6868417

74. Bohr T, Hjorth PG, Holst SC, Hrabětová S, Kiviniemi V, Lilius T, et al. The glymphatic system: Current understanding and modeling. iScience. 2022;25. doi:10.1016/j.isci.2022.104987

75. Waltl I, Kalinke U. Beneficial and detrimental functions of microglia during viral encephalitis. Trends in Neurosciences. 2022;45: 158–170. doi:10.1016/j.tins.2021.11.004

76. Li S-K, Leung RK-K, Guo H-X, Wei J-F, Wang J-H, Kwong K-T, et al. Detection and identification of plasma bacterial and viral elements in HIV/AIDS patients in comparison to healthy adults. Clin Microbiol Infect. 2012;18: 1126–1133. doi:10.1111/j.1469-0691.2011.03690.x

77. Li Y, Cao L, Ye M, Xu R, Chen X, Ma Y, et al. Plasma Virome Reveals Blooms and Transmission of Anellovirus in Intravenous Drug Users with HIV-1, HCV, and/or HBV Infections. Microbiol Spectr. 2022;10: e0144722. doi:10.1128/spectrum.01447-22

78. Liu K, Li Y, Xu R, Zhang Y, Zheng C, Wan Z, et al. HIV-1 Infection Alters the Viral Composition of Plasma in Men Who Have Sex with Men. mSphere. 2021;6: e00081–21. doi:10.1128/mSphere.00081-21

79. Li L, Deng X, Linsuwanon P, Bangsberg D, Bwana MB, Hunt P, et al. AIDS Alters the Commensal Plasma Virome. Journal of Virology. 2013;87: 10912–10915. doi:10.1128/JVI.01839-13

80. Guo Y, Huang X, Sun X, Yu Y, Wang Y, Zhang B, et al. The Underrated Salivary Virome of Men Who Have Sex With Men Infected With HIV. Front Immunol. 2021;12: 759253. doi:10.3389/fimmu.2021.759253

81. Bhagchandani T, Haque MMU, Sharma S, Malik MZ, Ray AK, Kaur US, et al. Plasma Virome of HIV-infected Subjects on Suppressive Antiretroviral Therapy Reveals Association of Differentially Abundant Viruses with Distinct T-cell Phenotypes and Inflammation. Curr Genomics. 2024;25: 105–119. doi:10.2174/0113892029279786240111052824

82. Ellis RJ, Marquine MJ, Kaul M, Fields JA, Schlachetzki JCM. Mechanisms underlying HIV-associated cognitive impairment and emerging therapies for its management. Nat Rev Neurol. 2023;19: 668–687. doi:10.1038/s41582-023-00879-y

83. Winston A, Spudich S. Cognitive disorders in people living with HIV. The Lancet HIV. 2020;7: e504–e513. doi:10.1016/S2352-3018(20)30107-7

84. Spudich S, Robertson KR, Bosch RJ, Gandhi RT, Cyktor JC, Mar H, et al. Persistent HIV-infected cells in cerebrospinal fluid are associated with poorer neurocognitive performance. J Clin Invest. 2019;129: 3339–3346. doi:10.1172/JCI127413

85. Chan P, Spudich S. HIV compartmentalization in the CNS and its impact in Treatment Outcomes and Cure Strategies. Curr HIV/AIDS Rep. 2022;19: 207–216. doi:10.1007/s11904-022-00605-1

86. Trunfio M, Atzori C, Pasquero M, Di Stefano A, Vai D, Nigra M, et al. Patterns of Cerebrospinal Fluid Alzheimer’s Dementia Biomarkers in People Living with HIV: Cross-Sectional Study on Associated Factors According to Viral Control, Neurological Confounders and Neurocognition. Viruses. 2022;14: 753. doi:10.3390/v14040753

87. Calcagno A, Celani L, Trunfio M, Orofino G, Imperiale D, Atzori C, et al. Alzheimer Dementia in People Living With HIV. Neurol Clin Pract. 2021;11: e627–e633. doi:10.1212/CPJ.0000000000001060

88. Bonnan M, Barroso B, Demasles S, Krim E, Marasescu R, Miquel M. Compartmentalized intrathecal immunoglobulin synthesis during HIV infection — A model of chronic CNS inflammation? J Neuroimmunol. 2015;285: 41–52. doi:10.1016/j.jneuroim.2015.05.015

89. De Almeida SM, Rotta I, Tang B, Vaida F, Letendre S, Ellis RJ. IgG intrathecal synthesis in HIV-associated neurocognitive disorder (HAND) according to the HIV-1 subtypes and pattern of HIV RNA in CNS and plasma compartments. J Neuroimmunol. 2021;355: 577542. doi:10.1016/j.jneuroim.2021.577542

90. Trunfio M, Mighetto L, Napoli L, Atzori C, Nigra M, Guastamacchia G, et al. Cerebrospinal Fluid CXCL13 as Candidate Biomarker of Intrathecal Immune Activation, IgG Synthesis and Neurocognitive Impairment in People with HIV. J Neuroimmune Pharmacol. 2023. doi:10.1007/s11481-023-10066-x

91. Yang E, Li MMH. All About the RNA: Interferon-Stimulated Genes That Interfere With Viral RNA Processes. Front Immunol. 2020;11: 605024. doi:10.3389/fimmu.2020.605024

92. Rehwinkel J, Gack MU. RIG-I-like receptors: their regulation and roles in RNA sensing. Nat Rev Immunol. 2020;20: 537–551. doi:10.1038/s41577-020-0288-3

93. Williams ME, Stein DJ, Joska JA, Naudé PJW. Cerebrospinal fluid immune markers and HIV-associated neurocognitive impairments: A systematic review. Journal of Neuroimmunology. 2021;358: 577649. doi:10.1016/j.jneuroim.2021.577649

94. Michetti F, Clementi ME, Di Liddo R, Valeriani F, Ria F, Rende M, et al. The S100B Protein: A Multifaceted Pathogenic Factor More Than a Biomarker. Int J Mol Sci. 2023;24: 9605. doi:10.3390/ijms24119605

95. Li S, Chen Y, Chen G. Cognitive disorders: Potential astrocyte-based mechanism. Brain Research Bulletin. 2025;220: 111181. doi:10.1016/j.brainresbull.2024.111181

96. Lee SY, Chung W-S. The roles of astrocytic phagocytosis in maintaining homeostasis of brains. Journal of Pharmacological Sciences. 2021;145: 223–227. doi:10.1016/j.jphs.2020.12.007

97. Jorgačevski J, Potokar M. Immune Functions of Astrocytes in Viral Neuroinfections. International Journal of Molecular Sciences. 2023;24: 3514. doi:10.3390/ijms24043514

98. Eimer WA, Vijaya Kumar DK, Navalpur Shanmugam NK, Rodriguez AS, Mitchell T, Washicosky KJ, et al. Alzheimer’s Disease-Associated β-Amyloid Is Rapidly Seeded by *Herpesviridae* to Protect against Brain Infection. Neuron. 2018;99: 56–63.e3. doi:10.1016/j.neuron.2018.06.030

99. Bourgade K, Le Page A, Bocti C, Witkowski JM, Dupuis G, Frost EH, et al. Protective Effect of Amyloid-β Peptides Against Herpes Simplex Virus-1 Infection in a Neuronal Cell Culture Model. J Alzheimers Dis. 2016;50: 1227–1241. doi:10.3233/JAD-150652

100. White MR, Kandel R, Tripathi S, Condon D, Qi L, Taubenberger J, et al. Alzheimer’s Associated β-Amyloid Protein Inhibits Influenza A Virus and Modulates Viral Interactions with Phagocytes. PLOS ONE. 2014;9: e101364. doi:10.1371/journal.pone.0101364

101. Galloway DA, Phillips AEM, Owen DRJ, Moore CS. Phagocytosis in the Brain: Homeostasis and Disease. Front Immunol. 2019;10. doi:10.3389/fimmu.2019.00790

102. Tang J, Zhang M, Liu N, Xue Y, Ren X, Huang Q, et al. The Association Between Glymphatic System Dysfunction and Cognitive Impairment in Cerebral Small Vessel Disease. Front Aging Neurosci. 2022;14: 916633. doi:10.3389/fnagi.2022.916633

103. Wang J, Zhou Y, Zhang K, Ran W, Zhu X, Zhong W, et al. Glymphatic function plays a protective role in ageing-related cognitive decline. Age Ageing. 2023;52: afad107. doi:10.1093/ageing/afad107

104. Veenhuis RT, Williams DW, Shirk EN, Abreu CM, Ferreira EA, Coughlin JM, et al. Higher circulating intermediate monocytes are associated with cognitive function in women with HIV. JCI Insight. 2021;6. doi:10.1172/jci.insight.146215

105. Rubin LH, Shirk EN, Pohlenz L, Romero H, Roti E, Dastgheyb RM, et al. Intact HIV reservoir in monocytes is associated with cognitive function in virally suppressed women with HIV. The Journal of Infectious Diseases. 2024; jiae460. doi:10.1093/infdis/jiae460

106. O’Flaherty BM, Li Y, Tao Y, Paden CR, Queen K, Zhang J, et al. Comprehensive viral enrichment enables sensitive respiratory virus genomic identification and analysis by next generation sequencing. Genome Res. 2018;28: 869–877. doi:10.1101/gr.226316.117

107. Motta I, Allice T, Romito A, Ferrara M, Ecclesia S, Imperiale D, et al. Cerebrospinal fluid viral load and neopterin in HIV-positive patients with undetectable viraemia. Antivir Ther. 2017;22: 539–543. doi:10.3851/IMP3140

108. Caligaris G, Trunfio M, Ghisetti V, Cusato J, Nigra M, Atzori C, et al. Blood-Brain Barrier Impairment in Patients Living with HIV: Predictors and Associated Biomarkers. Diagnostics (Basel). 2021;11: 867. doi:10.3390/diagnostics11050867

109. Blackstone K, Moore DJ, Franklin DR, Clifford DB, Collier AC, Marra CM, et al. Defining Neurocognitive Impairment in HIV: Deficit Scores versus Clinical Ratings. Clin Neuropsychol. 2012;26: 10.1080/13854046.2012.694479.

110. Beck AT, Steer RA, Brown G. Beck Depression Inventory–II. 2011. doi:10.1037/t00742-000

111. Petersen KJ, Yu X, Masters MC, Lobo JD, Lu T, Letendre S, et al. Sex-specific associations between plasma interleukin-6 and depression in persons with and without HIV. Brain Behav Immun Health. 2023;30: 100644. doi:10.1016/j.bbih.2023.100644

112. Hobkirk AL, Starosta AJ, De Leo JA, Marra CM, Heaton RK, Earleywine M. Psychometric Validation of the BDI-II Among HIV-Positive CHARTER Study Participants. Psychol Assess. 2015;27: 457–466. doi:10.1037/pas0000040

113. Chen S, Zhou Y, Chen Y, Gu J. fastp: an ultra-fast all-in-one FASTQ preprocessor. Bioinformatics. 2018;34: i884–i890. doi:10.1093/bioinformatics/bty560

114. Babraham Bioinformatics - FastQC A Quality Control tool for High Throughput Sequence Data. [cited 12 Jun 2023]. Available: https://www.bioinformatics.babraham.ac.uk/projects/fastqc/

115. Ewels P, Magnusson M, Lundin S, Käller M. MultiQC: summarize analysis results for multiple tools and samples in a single report. Bioinformatics. 2016;32: 3047–3048. doi:10.1093/bioinformatics/btw354

116. Wood DE, Lu J, Langmead B. Improved metagenomic analysis with Kraken 2. Genome Biol. 2019;20: 257. doi:10.1186/s13059-019-1891-0

117. Ondov BD, Bergman NH, Phillippy AM. Interactive metagenomic visualization in a Web browser. BMC Bioinformatics. 2011;12: 385. doi:10.1186/1471-2105-12-385

118. Lu J, Breitwieser FP, Thielen P, Salzberg SL. Bracken: estimating species abundance in metagenomics data. PeerJ Comput Sci. 2017;3: e104. doi:10.7717/peerj-cs.104

119. Madison JD, LaBumbard BC, Woodhams DC. Shotgun metagenomics captures more microbial diversity than targeted 16S rRNA gene sequencing for field specimens and preserved museum specimens. PLoS One. 2023;18: e0291540. doi:10.1371/journal.pone.0291540

120. Wright RJ, Comeau AM, Langille MGI. From defaults to databases: parameter and database choice dramatically impact the performance of metagenomic taxonomic classification tools. Microb Genom. 2023;9: mgen000949. doi:10.1099/mgen.0.000949

121. Nurk S, Meleshko D, Korobeynikov A, Pevzner PA. metaSPAdes: a new versatile metagenomic assembler. Genome Res. 2017;27: 824–834. doi:10.1101/gr.213959.116

122. Ansari MH, Ebrahimi M, Fattahi MR, Gardner MG, Safarpour AR, Faghihi MA, et al. Viral metagenomic analysis of fecal samples reveals an enteric virome signature in irritable bowel syndrome. BMC Microbiology. 2020;20: 123. doi:10.1186/s12866-020-01817-4

123. Wickham H. Data Analysis. In: Wickham H, editor. ggplot2: Elegant Graphics for Data Analysis. Cham: Springer International Publishing; 2016. pp. 189–201. doi:10.1007/978-3-319-24277-4_9

124. McDonald D, Clemente JC, Kuczynski J, Rideout JR, Stombaugh J, Wendel D, et al. The Biological Observation Matrix (BIOM) format or: how I learned to stop worrying and love the ome-ome. Gigascience. 2012;1: 7. doi:10.1186/2047-217X-1-7

125. Ma Y, Zhu L. A Review on Dimension Reduction. Int Stat Rev. 2013;81: 134–150. doi:10.1111/j.1751-5823.2012.00182.x

126. Murtagh F, Legendre P. Ward’s Hierarchical Agglomerative Clustering Method: Which Algorithms Implement Ward’s Criterion? J Classif. 2014;31: 274–295. doi:10.1007/s00357-014-9161-z

127. Milligan GW, Cooper MC. An examination of procedures for determining the number of clusters in a data set. Psychometrika. 1985;50: 159–179. doi:10.1007/BF02294245

128. Jiang R, Sun T, Song D, Li JJ. Statistics or biology: the zero-inflation controversy about scRNA-seq data. Genome Biology. 2022;23: 31. doi:10.1186/s13059-022-02601-5

